# Unraveling intratumoral heterogeneity through high-sensitivity single-cell mutational analysis and parallel RNA-sequencing

**DOI:** 10.1101/474734

**Authors:** Alba Rodriguez-Meira, Gemma Buck, Sally-Ann Clark, Benjamin J Povinelli, Veronica Alcolea, Eleni Louka, Simon McGowan, Angela Hamblin, Nikolaos Sousos, Nikolaos Barkas, Alice Giustacchini, Bethan Psaila, Sten Eirik W Jacobsen, Supat Thongjuea, Adam J Mead

## Abstract

Single-cell RNA-sequencing has emerged as a powerful tool to resolve transcriptional heterogeneity. However, its application to study cancerous tissues is currently hampered by the lack of coverage across key mutation hotspots in the vast majority of cells, which prevents correlation of genetic and transcriptional readouts from the same single cell. To overcome this, we developed TARGET-seq, a method for the high-sensitivity detection of multiple mutations within single-cells from both genomic and coding DNA, in parallel with unbiased, high-depth whole transcriptome analysis. We demonstrate how this technique uniquely resolves transcriptional and genetic tumor heterogeneity in myeloproliferative neoplasm stem/progenitor cells, providing insights into deregulated pathways of mutant and non-mutant cells. TARGET-seq provides a powerful tool to resolve molecular signatures of genetically distinct subclones of tumor cells.

## INTRODUCTION

Resolving intratumoral heterogeneity (ITH) is critical for our understanding of tumor evolution and resistance to therapies, which in turn is required for the development of effective cancer treatments and identification of biomarkers for precision medicine (Alizadeh et al., 2015; McGranahan and Swanton, 2017). The best characterized source of ITH is at the genetic level, where advances in next generation sequencing (NGS) techniques at the bulk and single-cell level have described the genetic diversity within tumors with unprecedented resolution (Kandoth et al., 2013; Vogelstein et al., 2013). However, a number of other factors beyond somatic mutations contribute to ITH. For example, some tumors are hierarchically organised, including the presence of cancer stem cells (CSC), which propagate disease relapse. The genetic events underlying tumor evolution originate in CSCs, which in some tumors, are rare within the total tumor bulk population (Clevers, 2011; Magee et al., 2012; Woll et al., 2014). Furthermore, the normal cellular counterparts of CSCs, which lack genetic mutations, can be difficult to distinguish from malignant cells as they may share multiple phenotypic features, but can nevertheless be informative for disease biology (Giustacchini et al., 2017). Consequently, resolving ITH requires methods that allow these multiple layers of heterogeneity to be teased apart.

To better understand the functional consequences of ITH, a potentially powerful approach is to link genetic ITH with transcriptional signatures of distinct subpopulations of tumour cells. A number of studies have begun to apply single-cell RNA-sequencing (scRNA-seq) to characterize the cellular landscape of a number of different malignancies (Patel et al., 2014; Tirosh et al., 2016a; Tirosh et al., 2016b; Venteicher et al., 2017). These pioneering studies have demonstrated the power of scRNA-seq to identify the different cell types encompassed within a tumor (Patel et al., 2014; Tirosh et al., 2016a; Tirosh et al., 2016b), including cells with “stemness” signatures (Tirosh et al., 2016b) and characterisation of developmental hierarchies of tumor cells (Tirosh et al., 2016b). However, although scRNA-seq approaches can readily resolve such transcriptional heterogeneity, current techniques do not allow parallel mutational analysis due to lack of coverage across mutation hotspots (Kiselev et al., 2017; Patel et al., 2014; Tirosh et al., 2016b). This integration of mutation and transcriptional information is crucial in order to link genetic evolution events to the cell of origin, of considerable importance as serial mutation acquisition might occur within distinct and developmentally ordered stem/progenitor cell types, as described in acute leukemia (Jan et al., 2012). Furthermore, mutation analysis would not only be important for the identification of mutant cells, but also to unravel disrupted gene expression in non-mutant cells, which may be cell-extrinsically mediated and of clinical importance (Giustacchini et al., 2017). In order to overcome this current limitation in single cell genomic techniques, we set out to develop a method that would allow combined scRNA-seq with high sensitivity mutation analysis.

## DESIGN

The limitation of current scRNA-sequencing techniques for the detection of mutations in single cells partly relates to amplification of only the 3’ or 5’ region of transcripts with commonly used “end-counting” scRNA-seq techniques (Hedlund and Deng, 2018). Consequently, most mutations within the body of a gene are not covered by sequencing reads. However, scRNA-seq techniques that amplify full-length transcripts, such as Smart-seq2 (Hedlund and Deng, 2018; Picelli et al., 2013), also have very poor sensitivity to detect expression of most genes in most cells (Figure 1a and 1b), which precludes high sensitivity mutational analysis. Furthermore, the vast majority of mutations identified in cancer are single nucleotide variants (SNV) and small indels, which may be either heterozygous or associated with loss of heterozygosity (LOH) (Vogelstein et al., 2013), with important functional consequences (Kharazi et al., 2011). Therefore, a key challenge in the field is to minimize allelic dropouts (ADO), in order to ensure the detection of both alleles from a single-cell.

**Figure 1.**
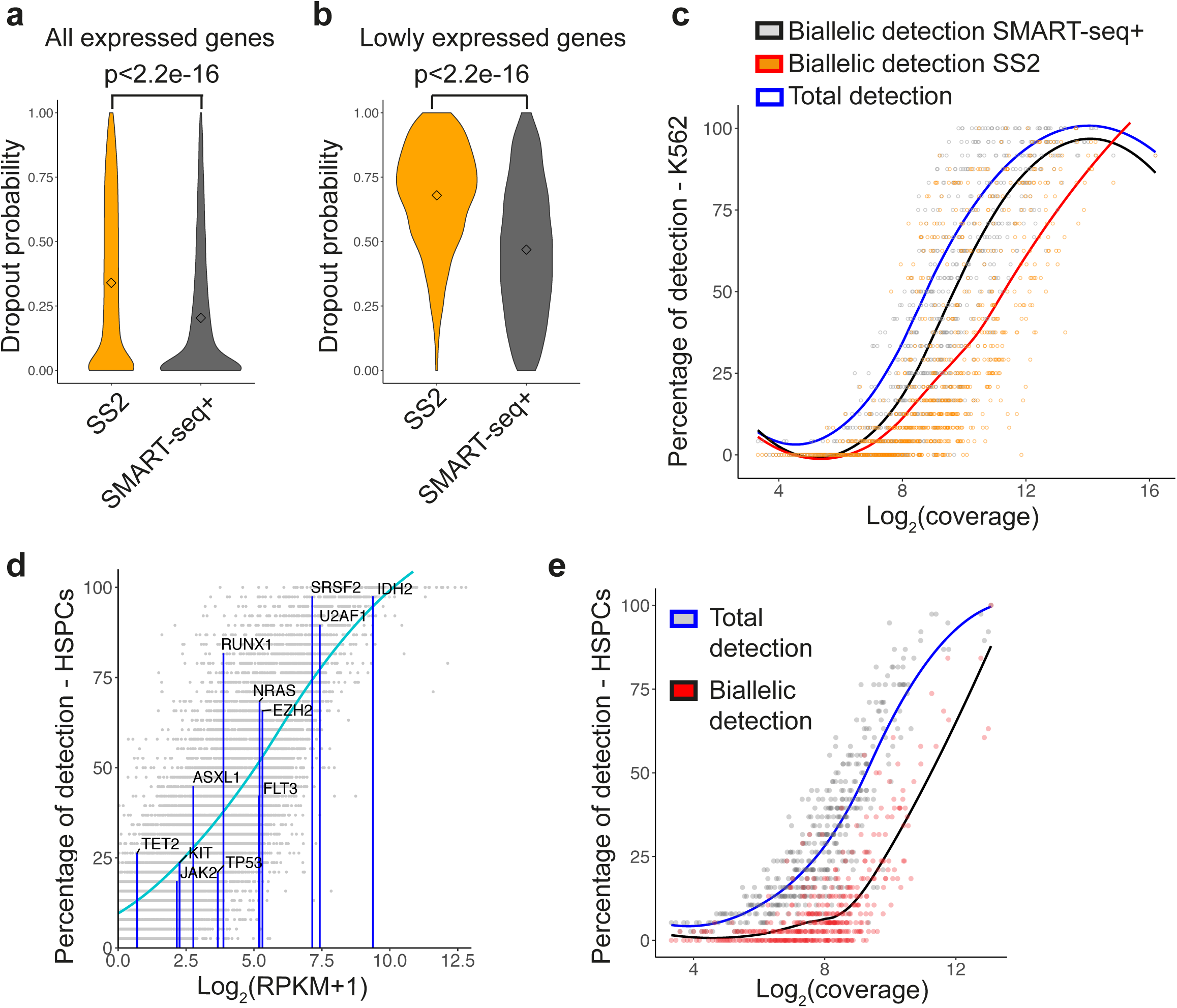
Single-cell RNA-sequencing is associated with high levels of allelic dropout. **(a)** Comparison of dropout frequency between SMART-seq2 and SMART-seq+ methods (n=48 single K562 cells) for all genes expressed in K562 bulk samples (RPKM>1; 9096 genes). P-value from two-tailed unpaired Student t test is shown on the top of the graph. Points represent the mean for each group. **(b)** Comparison of dropout frequency for lowly expressed genes (2>RPKM>1; 1347 genes) between SMART-seq2 and SMART-seq+ methods (n=48 single K562 cells). P-value from two-tailed unpaired Student t test is shown on the top of the graph. Points represent the mean for each group.**(c)** Percentage of total (dark blue line) or bi-allelic detection in heterozygous SNVs for Smart-seq2 (orange dots and red line) or optimized SMART-seq+ (grey dots and black line) chemistries (n=48 single K562 cells). Lines represent the mean percentage of detection (y-axis) with respect to coverage (x-axis) and points represent individual SNVs. **(d)** Total percentage of detection of selected myeloid genes in Lin-CD34+CD38-hematopoietic stem/progenitor cells (HSPC; n=38; y-axis) with respect to the average level of expression for each gene (log2(RPKM+1); x-axis). Blue bars represent detection of specific gene transcripts that are mutated in myeloid malignancies. The blue line represents the average percentage of detection for a certain expression value, and each grey dot represents an individual transcript. **(e)** Total versus bi-allelic percentage of detection of heterozygous SNVs in the same single cells as in (d) with respect to the total number of reads spanning that position (log2(coverage); x-axis). The blue line and grey points represent the total percentage of detection for a certain heterozygous position. The black line and red points indicate the detection of both alleles (at least 5% of reads mapping to either of the alleles).

It remains unclear whether the high ADO rates and lack of coverage across mutation hotspots with scRNA-seq data is primarily due to technical dropouts due to inefficient reverse transcription (RT) and/or PCR amplification, or true biological heterogeneity in expression of mutant transcripts across single cells. We therefore first optimized the Smart-seq2 reverse transcription and amplification enzymatic conditions (SMART-seq+; Table S1), resulting in a significant reduction in dropout rates (Figure 1a), particularly for lowly expressed genes (Figure 1b), a 25 % increase in the number of genes detected per cell (Figure S1a) and a reduction in library bias (Figure S1b). However, despite the improved sensitivity for detection of gene expression with SMART-seq+, ADO rates remained exceedingly high for most genes (Figure 1c), which currently precludes reliable mutational analysis using scRNA-seq (Povinelli et al., 2018). We therefore concluded that, due to the stochastic nature of gene expression in single cells, improving the sensitivity for the analysis of coding DNA (cDNA) alone is unlikely to provide sufficient sensitivity for detection of most cancer-associated mutations at the single cell level.

To overcome this problem, it is required to detect mutations from genomic DNA (gDNA) in parallel with cDNA. Techniques to study gDNA and mRNA from the same single cell have been previously described (Dey et al., 2015; Hou et al., 2016; Macaulay et al., 2015). However, these techniques either require physical separation of gDNA and mRNA (Han et al., 2018; Hou et al., 2016; Macaulay et al., 2015), inevitably resulting in some loss of genetic material and consequently limiting their sensitivity, or rely on the parallel amplification of total gDNA and mRNA with subsequent masking of coding regions (Dey et al., 2015). These technical constraints restrict the sensitivity of such techniques for the confident detection of specific point mutations. Whole genome amplification also introduces significant expense to the method, and has inherently high allelic dropout and false positive rates (Hosokawa et al., 2017; Wang et al., 2014). As a result, up to now, these techniques have not been widely used for parallel mutation and scRNA-seq analysis in cancer. Methods that combine targeted single cell gene expression and mutation analysis have also been reported (Cheow et al., 2016; Wang et al., 2017), but these approaches have the limitation that only the expression of a limited number of pre-selected genes can be analyzed per cell.

Recently, we have described a novel method for the high sensitivity detection of BCR-ABL transcripts in parallel with scRNA-seq, which we applied to characterize the molecular signatures of stem cells in chronic myeloid leukemia (Giustacchini et al., 2017). Whilst this study highlights the potential power of linking mutation and transcriptome information in single-cells, this method is also dependent on the expression of the targeted gene/allele in all mutated cells. This approach was effective in the specific case of BCR-ABL fusion gene, but for many autosomal genes, expression is undetectable or highly allelic biased (Deng et al., 2016) in the majority of transcriptionally active and highly proliferative K562 cells (Figure 1c) and also in quiescent Lin^-^CD34^+^CD38^-^ primary human hematopoietic stem/progenitor cells (HSPCs; Figures 1d and 1e), which makes this method unsuitable to profile most mutations found in cancer. Moreover, this approach precludes analysis of non-coding mutations, with key roles in tumorigenesis (Khurana et al., 2016). We therefore developed a method, named TARGET-seq, which dramatically reduces ADO and also enables efficient detection of non-coding mutations by allowing parallel, targeted mutation analysis of gDNA and cDNA alongside scRNA-seq from the same single-cell.

## RESULTS

### TARGET-seq dramatically increases the sensitivity of mutation detection in single cells

In order to improve detection of specific mRNA and gDNA amplicons, we extensively modified the SMART-seq2 protocol (Hedlund and Deng, 2018; Picelli et al., 2013). To improve release of gDNA, we modified the lysis procedure to include a mild protease digestion (Figure 2a; See Detailed Protocol), which was subsequently heat-inactivated to avoid inhibiting RT and PCR steps. Target-specific primers for cDNA and gDNA were added to cDNA synthesis and PCR amplification steps, which also used modified enzymes (Figure 2a, Table S2) providing a more efficient RT and PCR amplification. An aliquot of the pre-amplified gDNA/cDNA libraries was used for targeted NGS of specific cDNA and gDNA amplicons, and another aliquot, for whole transcriptome library preparation. The targeted mutation analysis and scRNA-seq libraries are sequenced and analysed independently.

**Figure 2.**
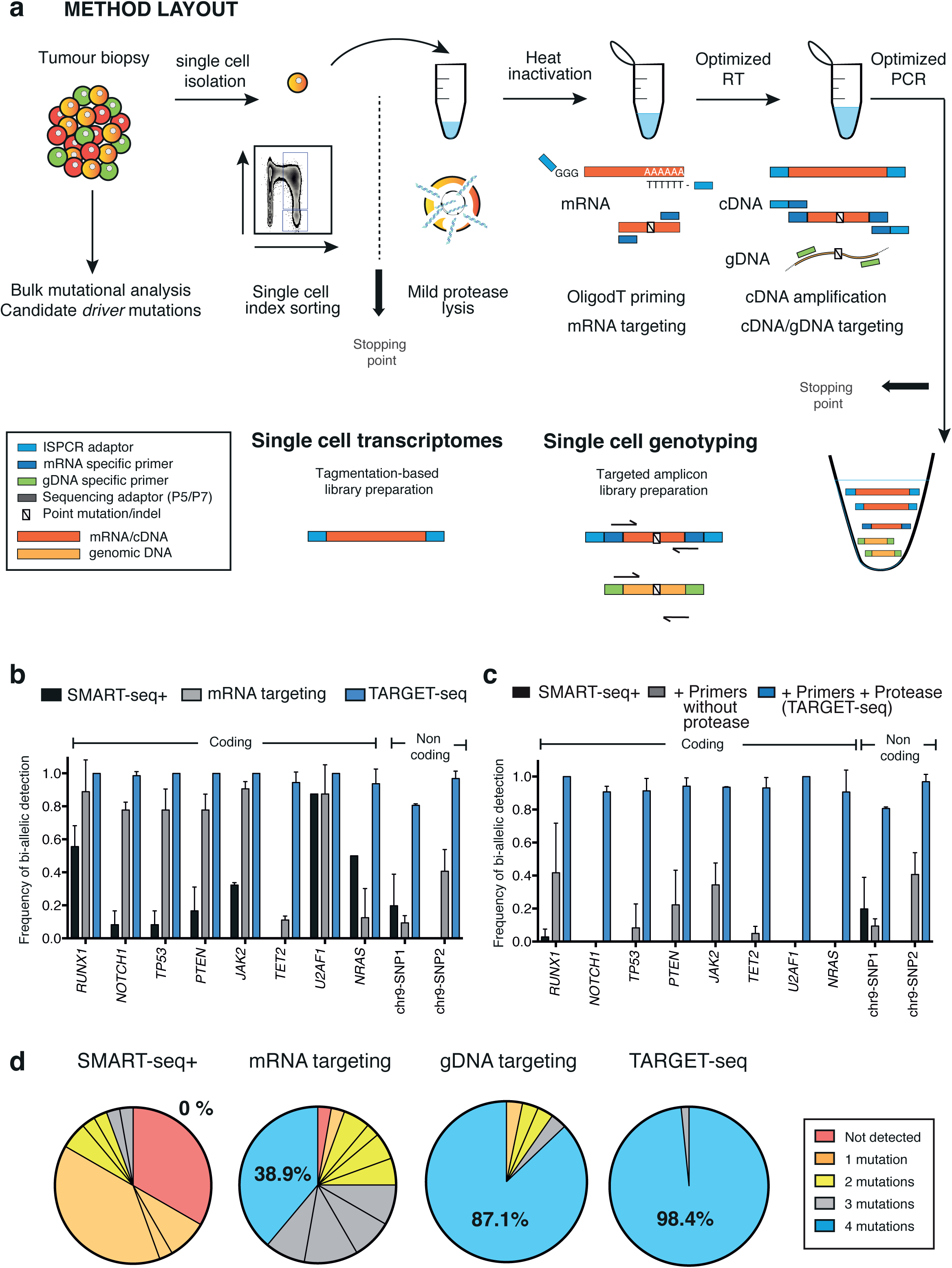
TARGET-seq: a method for high sensitivity mutational detection and parallel whole transcriptome from the same single cell. **(a)** Schematic representation of the method. First, single cells are isolated by fluorescent-activated cell sorting (FACS) into 96 well plates containing TARGET-seq lysis buffer. Plates are then incubated at 72 C in a thermocycler to inactivate the protease contained within the lysis buffer. Subsequently, retrotranscription mix is added to the plates, in which OligodT-ISPCR is used to prime total polyadenylated mRNA molecules and target-specific primers are used to prime molecules of interest. In a subsequent PCR step, ISPCR primers are used to amplify polyadenylated cDNA and target specific primers are used to amplify cDNA and gDNA molecules of interest. **(b)** Frequency of detection of heterozygous mutations using TARGET-seq in ten coding and non-coding regions, as compared to SMART-seq+ and mRNA targeting approaches (n=376 cells, 2-3 independent experiments per amplicon; bar graph represents mean±sd). **(c)** Frequency of detection of heterozygous mutations for the ten validated amplicons from (b), showing exclusively results from targeted genomic DNA sequencing. Bar graph represents mean±sd.**(d)** Frequency of detection of heterozygous mutations in JURKAT cells with SMART-seq+ (n=36 cells), mRNA targeting (n=36 cells), gDNA targeting (n=62 cells) and TARGET-seq (n=62 cells) when profiling four different mutations in the same single cell (*RUNX1, NOTCH1, PTEN* and *TP53*) in three independent experiments. Each slice of the pie chart represents a different combination of mutations and each color represents the number of mutations detected per single cell.

TARGET-seq dramatically improved detection of ten mutation hotspots in clonal cell lines, including SNV and small indels across both coding and non-coding regions (Figure 2b). Notably, gDNA amplicons alone achieved a mean 93 percent biallelic mutation/SNV detection (Figure 2c; schemes of the variant calling pipeline and specific examples of variant calling can be found in Figure S2a and Figure S2b, respectively). Importantly, mutational analysis from raw RNA-sequencing reads, as opposed to targeted re-sequencing, was not possible in almost all cells due to lack of coverage (Figure S2c), despite reaching a mean sequencing depth of 2.93 million reads/cell.

We next tested whether TARGET-seq would improve detection of combinations of mutations in single-cells. We profiled four different mutations in a clonal T-cell leukemia diploid cell line carrying heterozygous mutations in *NOTCH1, RUNX1, TP53* and *PTEN* (JURKAT cells). Using SMART-seq+, all 4 mutations were detected in none of the cells analyzed, compared with 38.9% by mRNA targeting, 87.1% by gDNA targeting and 98.4% by TARGET-seq (combined mRNA+gDNA targeting; Figure 2d). Therefore, TARGET-seq provides extremely high sensitivity for the detection of multiple mutations in the same single-cell, which is essential for reliable reconstruction of tumor phylogenetic trees.

### TARGET-seq produces unbiased transcriptomic readouts from single cells

To determine whether TARGET-seq introduces a bias in the single cell whole transcriptome data, we evaluated its performance in two cell lines (JURKAT and SET2) and in primary human HSPCs. Cells clustered by cell type and not by method (Figure 3a, 3b), with no significant differences in the number of genes detected between methods (Figure 3c). Sequencing quality controls (QC; Figure S3a), numbers of cells passing QC (Figure S3b) and transcript coverage (Figure S3c) was comparable between SMART-seq+ and TARGET-seq, with good correlations of gene expression, including genes selected for targeted amplification (Figure 3d, Figures S3d, S3e). Similarly, ERCC spike-in controls revealed high correlations between methods (Figure 3e, Figures S3f, S3g) and cDNA traces were comparable (Figures S3h, S3i and S3j). These results demonstrate that TARGET-seq allows accurate mutation detection with parallel unbiased scRNA-seq of the same single-cell.

**Figure 3.**
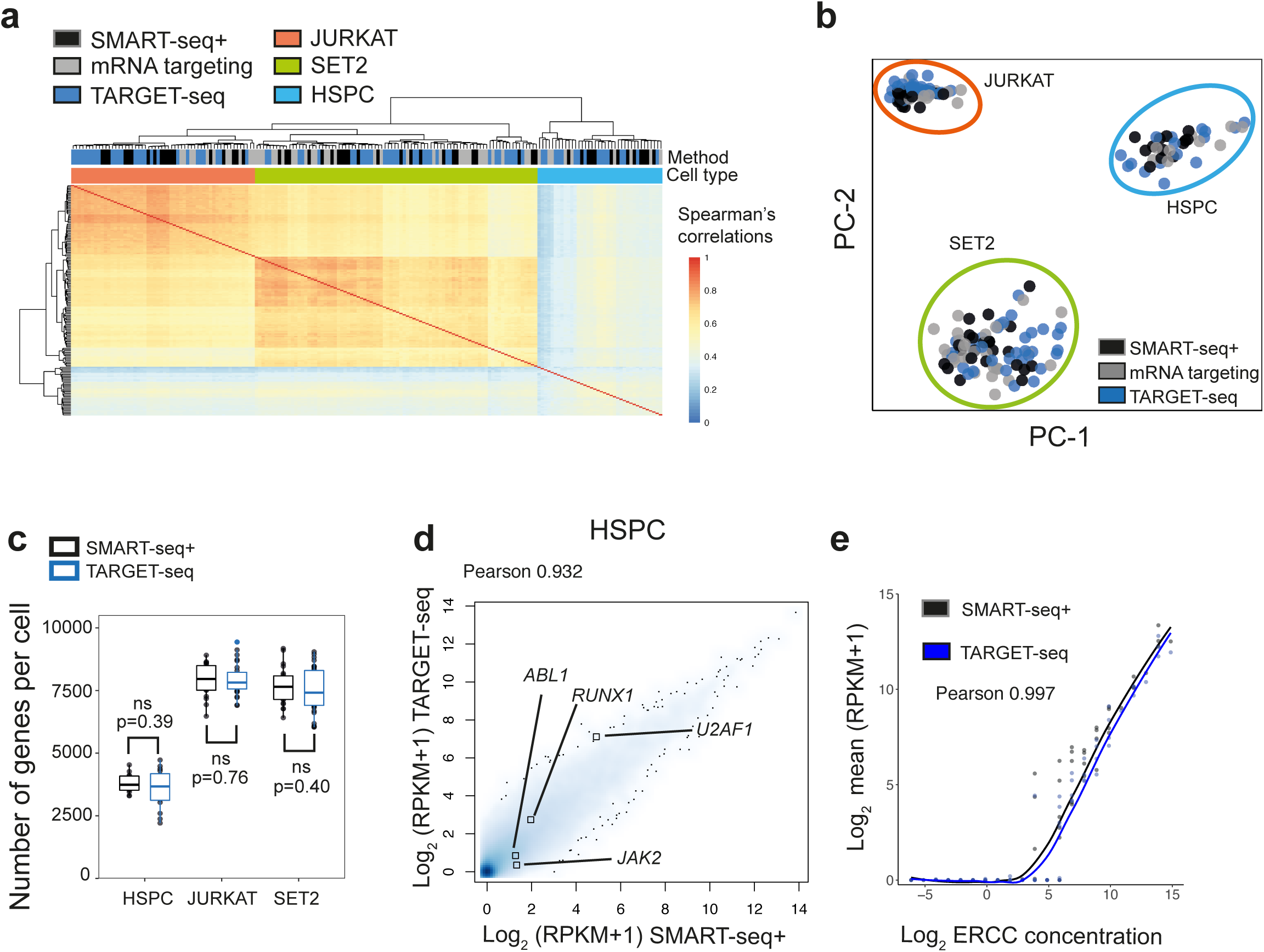
Unbiased whole transcriptome analysis of single cells using TARGET-seq. **(a)** Unsupervised hierarchical clustering of Spearman’s correlations from 180 single cells (JURKAT, n=56; SET2, n=86 and HSPC, n=38) using 15672 expressed genes (RPKM>=1). **(b)** Principal component analysis of HSPCs, SET2 and JURKAT cells from (a) using SMART-seq+, mRNA targeting or TARGET-seq. **(c)** Number of detected genes per cell (RPKM≥1) in HSPCs, SET2 and JURKAT cell lines using SMART-seq+ or TARGET-seq. **(d)** Whole transcriptome Pearson’s correlation between SMART-seq+ and TARGET-seq ensembles (mean RPKM values per condition) in HSPCs. The expression values for the genes targeted are highlighted. **(e)** Pearson’s correlation between mean ERCC expression values from SMART-seq+ and TARGET-seq in HSPCs.

### The stem cell compartment of MPN patients is genetically and transcriptionally heterogeneous

We next applied TARGET-seq to analyze 458 HSPCs from five patient samples with myeloproliferative neoplasms (MPN), carrying different combinations of *JAK2V617F, EZH2* and *TET2* mutations (Table S3). Two normal donors were also included as controls (Table S3). We isolated Lin-CD34+ cells by FACS (Figure S4) and indexed the cells for CD38, CD90, CD45RA and CD123, allowing assessment of clonal involvement in different stem-progenitor cell compartments (Majeti et al., 2007). All mutations identified in total mononuclear cells (MNCs; Figure S5a) were also detected in single-cells within the Lin-CD34+ compartment using TARGET-seq, which was superior to mRNA targeting alone (Figure S5b and S5c), and revealed subclonal mutations in all patients analysed with striking inter-patient heterogeneity in subclonal architecture. This allowed the order of acquisition of mutations to be determined (Figure S5c), of importance for MPN biology (Ortmann et al., 2015). For example, in patient SMD32316, we could determine that a *TET2* mutation was acquired after the *JAK2V617F* mutation, whereas in patient OX2123, *TET2* mutation was acquired before *JAK2V617F*. In two patients with similar *JAK2V617F* VAF in bulk MNCs, the low percentage of allelic dropout achieved with TARGET-seq analysis of single cells, revealed that *JAK2V617F* was heterozygous in most Lin-CD34+CD38-cells in one of these patients, with a normal distribution within the different Lin-CD34+CD38-stem-progenitor fractions (Figure 4a), whereas in the other patient a lower fraction of clonally involved Lin-CD34+CD38-cells were homozygous, and predominantly with a CD90+CD45RA+ aberrant phenotype (Figure 4b), also reported in other myeloid malignancies (Dimitriou et al., 2016). The ability to reliably distinguish heterozygous versus homozygous JAK2V617F mutation is of considerable importance for MPN biology (Li et al., 2014), and also more broadly in cancer as mutant allele specific imbalance is common in malignant disease, particularly during disease progression (Soh et al., 2009).

**Figure 4.**
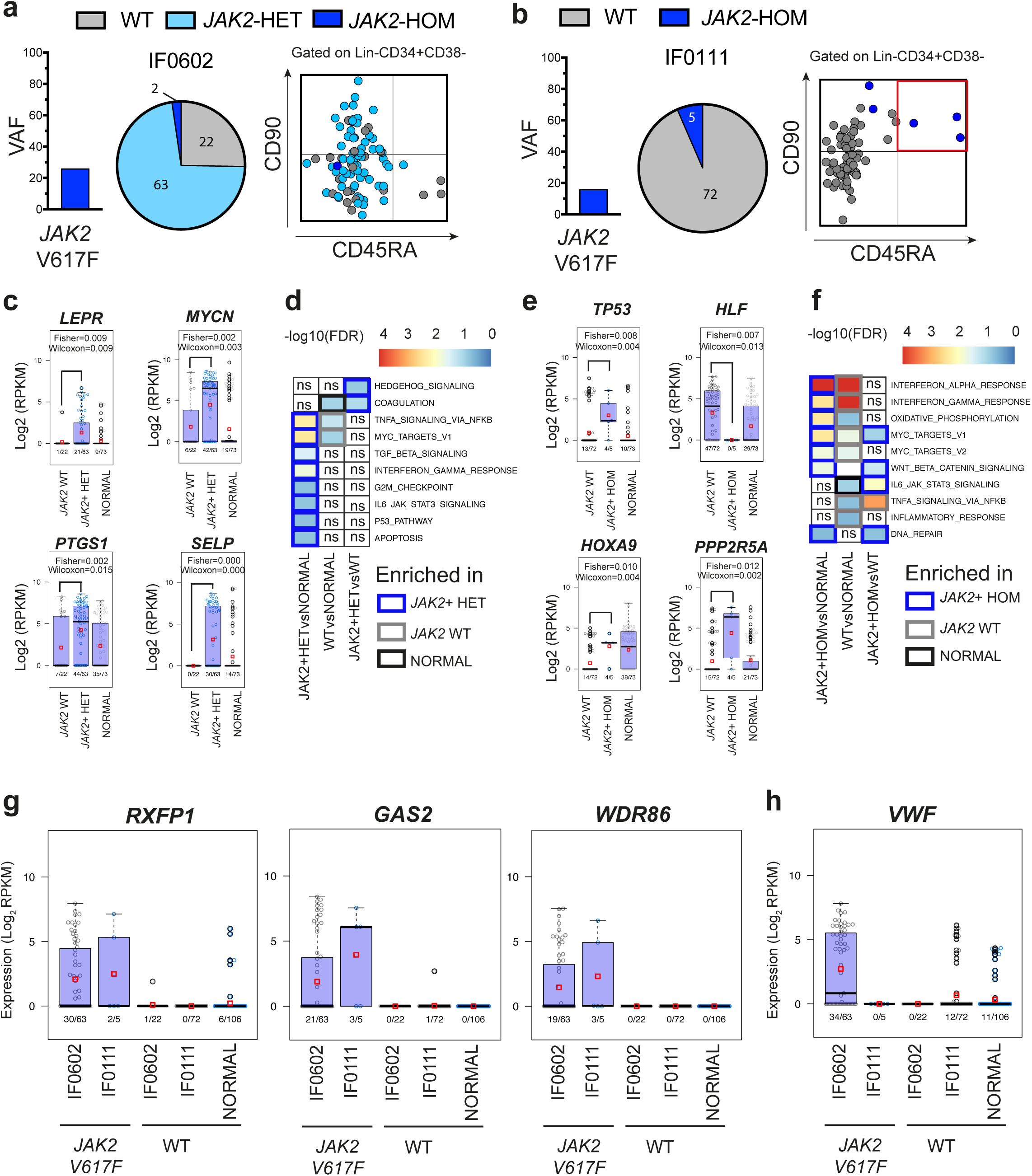
TARGET-seq reveals genetic and transcriptional heterogeneity in the stem cell compartment of MPN patients. **(a, b)** Variant allele frequency of *JAK2V617F* mutation (left), as identified by traditional bulk sequencing of total MNCs, proportion of single cells that carry the mutation in the Lin-CD34+CD38-compartment (center) and integration of index sorting with mutational information (right) for patients IF0602 **(a)** and IF0111 **(b)**. **(c-f)** Analysis of disrupted gene expression associated with *JAK2V617F* in HSPCs. Beeswarm plots of selected differentially expressed genes between **(c)** *JAK2* WT and *JAK2V617F* heterozygous mutant cells from patient IF0602 or **(e)** *JAK2* WT and *JAK2V617F* homozygous mutant cells from patient IF0111. Expression values for single cells from two normal donors (‘NORMAL’) are also shown. Each dot represents the expression value for each single cell; red squares represent mean expression values for each group and boxes represent median and quartiles. Fisher’s test and Wilcoxon test p-values are shown on the top each graph; expression frequencies are shown on the bottom of each bar for each group. Table S5a (patient IF0602) and Table S5c (patient IF0111) show all significant differentially expressed genes. **(d)** GSEA analysis of *JAK2* WT and *JAK2V617F* heterozygous mutant cells from patient IF0602 or **(f)** *JAK2* WT and *JAK2V617F* homozygous mutant cells from patient IF0111 as well as cells from normal donors (‘NORMAL’). Heatmap is colored according to –log10(FDR q-values) for each comparison, using a FDR q-value cut-off <0.25. The white color with ‘ns’ represents non-significance. The borders of each square of the heatmap are colored according to the experimental group in which a particular pathway is enriched (grey borders, enriched in *JAK2* WT cells; blue borders, enriched in *JAK2V617F* mutant cells; black borders, enriched in cells from normal donors). Table S5b (patient IF0602) and Table S5d (patient IF0111) show results for all genesets tested with FDR q-value<0.25. **(g)** Beeswarm plots of selected genes (identified as biomarkers of *JAK2* mutant cells independently of the patient analyzed) expression values across HSPCs from patients IF0602, IF0111 (*JAK2* WT and *JAK2V617F* mutant cells) and two normal donors (‘NORMAL’). Expression values and frequencies for *JAK2* WT and *JAK2V617F* mutant cells for each patient is shown, as well as cells from normal donors (‘NORMAL’). **(h)** Beeswarm plot of *VWF* expression values across HSPCs from patients IF0602, IF0111 (*JAK2* WT and *JAK2* V617F mutant cells) and two normal donors (‘NORMAL’). Only cells passing both genotyping and RNA-seq QC are shown for each patient.

TARGET-seq analysis uniquely allowed wild-type HSPCs to be reliably distinguished from *JAK2V617F* mutant cells in the same samples, revealing aberrant expression of biologically relevant genes such as *LEPR* (Jiang et al., 2008) and oncogenes such as *MYCN, TP53* or *PPP2R5A,* as well as biologically relevant pathways in heterozygous (Figure 4c,4d) and homozygous (Figure 4e,4f) *JAK2V617F* mutated HSPCs, including upregulation of hedgehog (Figure 4d) and Wnt/beta-catenin (Figure 4f) pathway associated transcription (Table S5). Importantly, this analysis allowed us to identify candidate biomarkers for *JAK2V617F* mutations in HSPCs (Figure 4g), two of which have been previously identified as candidate biomarkers in chronic myeloid leukemia (*RXFP1, GAS2*) (Giustacchini et al., 2017), and a novel candidate gene, *WDR86*, belonging to the WD-repeat protein family. Interestingly, *VWF*, a marker of platelet-biased stem cells (Sanjuan-Pla et al., 2013), was specifically upregulated in *JAK2V617F* heterozygous mutant cells from patient IF0602, whose disease was characterized by abnormal megakaryocytic differentiation and myelofibrosis, but not in *JAK2V617F* homozygous mutant cells from patient IF0111, who had a polycythemia phenotype (Figure 4h). These data support that transcriptional lineage priming in the HSPC compartment might be linked to the disease phenotype in MPN.

Crucially, *JAK2V617F* mutated HSPC could not be identified computationally by dimensionality reduction methods or hierarchical clustering (Figure S6a-d), which was not affected by cell cycle phase (Figure S6e, S6f). *JAK2V617F* could not be identified using a recently published single-cell k-means clustering method (Kiselev et al., 2017), SC3, previously used to specifically distinguish genetically-distinct subclones of cells (Figure S6g). *JAK2V617F* mutant and wild-type cells from the same patient could only be identified by hierarchical clustering using differentially expressed genes between mutant and wild-type cells (Figure S6h), the identification of which was only possible using TARGET-seq.

### Distinct genetic subclones present unique transcriptional signatures

TARGET-seq also uniquely allowed global transcriptome comparison of wild-type cells from patient samples and normal controls. Intriguingly, this analysis established that wild-type HSPCs from patients with MPN were transcriptionally distinct from normal donor HSPCs (Figure 5a), with enrichment of inflammatory pathways associated with TNF-alpha and IFN signalling (Figure 4d, Figure 5b). These results may indicate effects of the MPN microenvironment on the wild-type cells from the same patient, with a similar finding demonstrated to have clinically predictive value in chronic myeloid leukemia (Giustacchini et al., 2017). Interestingly, *JAK2V617F* mutant as well as WT HSPCs from patient IF0111 showed strong IFN signaling signatures, corresponding to the ongoing treatment of this particular patient with interferon, thus providing an additional layer of validation of the transcriptional signatures obtained (Figure 4f, Figure 5b).

**Figure 5.**
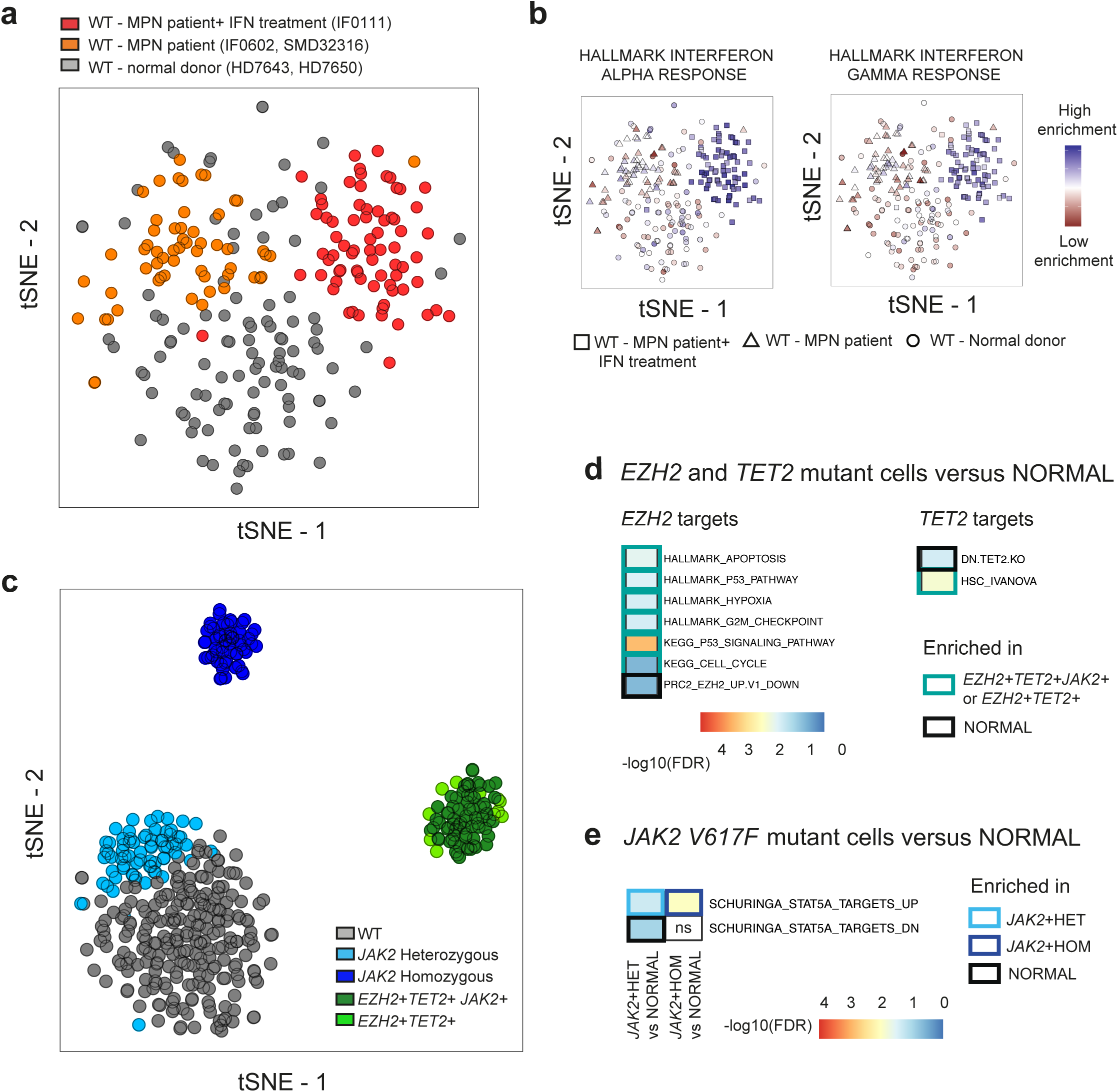
TARGET-seq reveals distinct transcriptional signatures associated with the presence or absence of somatic mutations in single HSPCs. **(a)** tSNE representation of 236 WT cells from the three patient samples in which WT cells are present (IF0602, SMD32316, IF0111), and cells from two normal donors (HD7650, HD7643), using 5365 highly variable genes. Cells from normal donors are colored in grey, cells from MPN patients are colored in orange (SMD32316, IF0602) or red (IF0111; patient being treated with interferon). **(b)** Enrichment of IFN-α (left panel) or IFN-γ (right panel) signalling gene signatures in WT cells from 3 patient samples and two normal donors, as a projection of ssGSEA results using the same tSNE coordinates from the cells of the specific donors/patients shown in (a). Each shape represents a group of donors. **(c)** tSNE representation of 448 cells from five patient samples and two normal controls, using the top 2000 genes as measured by the Gini index from the random forest analysis. Only genotypes present in at least five cells were analyzed. Gene expression matrix was batch and donor corrected using limma, whilst preserving genotypes. **(d-e)** Enrichment of *EZH2* related pathways, *TET2* related pathways (**d**) or JAK/STAT pathway (**e)** in cells carrying mutations in these genes compared to (n=106) single cells from two normal donors. Heatmap is colored according to –log10(FDR q-values) for each comparison, using a FDR q-value cut-off <0.25. A complete list of all significant genesets can be found in Table S6.

Analysis of combinations of mutations using the top 2000 important genes from random forest analysis (Figure 5c) showed striking clustering of HSPCs of the same genotype from multiple different patients. HSPCs carrying mutations in epigenetic modifiers had a highly distinct transcriptomic signature whereas cells carrying only *JAK2V617F* mutations more closely resembled the transcriptome of wild-type cells (Figure 5c). *EZH2* mutant cells showed enrichment in expression of PRC2 targets, and pathways previously identified to be correlated with *EZH2* mutations such as hypoxia, cell cycle, apoptosis and P53 signalling (Figure 5d; Table S6). *TET2* mutant cells also showed enrichment in HSC-related genes and a negative enrichment in genes downregulated upon *TET2* knock-out (Zhang et al., 2016) (Figure 5d, Table S6). Moreover, *JAK2V617F* cells showed dysregulation of *STAT5A* targets in a gene-dosage dependent manner, with JAK2V617F homozygous cells presenting a higher enrichment in *STAT5A* targets than *JAK2V617F* heterozygous cells (Figure 5e, Table S6). Taken together, these data demonstrate that TARGET-seq allows distinct and biologically relevant molecular signatures of HSPC subclones in MPN to be unravelled, representing a powerful tool for biomarker and therapeutic target discovery.

## DISCUSSION

With the advent of molecularly targeted therapy in cancer (Bible and Ryder, 2016; Longo, 2017; Luke et al., 2017), clinical remissions and clonal responses can be readily achieved in many patients. However, relapse frequently occurs, often associated with evidence of clonal evolution, likely reflecting ITH already present at diagnosis (Smith et al., 2017), with differential response to the targeted therapy in distinct tumor subclones. Therefore, to understand mechanisms of disease resistance, it is crucial to resolve clonal heterogeneity of tumors, and dissect the molecular signatures of responsive and resistant subclones of tumor cells. Whilst scRNA-seq offers great potential to resolve transcriptomic signatures of tumor subclones, up to now it has not been possible to correlate scRNA-seq data with mutation analysis due to lack of coverage in the scRNA-seq reads for small indels or point mutations, although large chromosomal aberrations can be detected (Tirosh et al., 2016a). For example, in an analysis of 430 cells from five primary glioblastomas using SMARTer Kit (v1), detection of *TP53* point mutations was possible in only 3 cells, leading the authors to conclude that “heterogeneity generated by focal alterations and point mutations will be grossly underappreciated using this method” (Patel et al., 2014). In two more recent study of gliomas, from 22 mutations analyzed, reads spanning the position of the mutations were detected in 0.4 to 8.7 % of the cells in the first study (Tirosh et al., 2016b), whereas scRNA-seq reads covered the exact site of the *IDH1* mutation in <40% of the cells, achieving detection of both the wild-type and the mutant alleles only in ∼5% of the malignant cells in the second study, reflecting the limited sensitivity for mutation detection (Venteicher et al., 2017). Although methods for the parallel sequencing of the whole-transcriptome and whole-genome of single-cells have previously been reported, these methods are not well suited to high-sensitivity mutation detection due to high ADO rates (Dey et al., 2015; Macaulay et al., 2015). Furthermore, these approaches are relatively costly due to the requirement for whole genome amplification. Consequently, up to now, such techniques have not been widely used for tumor analysis.

We herein report a novel single-cell RNA-sequencing and genotyping method that overcomes current limitations in available RNA-sequencing techniques by providing a simple, easily implemented and customizable protocol for high-sensitivity mutation detection with parallel high-depth, unbiased whole transcriptome analysis. TARGET-seq has clear advantages above other available scRNA-seq methodologies, providing improved complexity of scRNA-seq libraries and dramatically improved ability to detect multiple mutations in the same single cell, primarily attributable to the detection of gDNA through modified cell lysis and high sensitivity targeted amplification. The high sensitivity for biallelic detection of mutations provided by our technique is also of considerable importance as loss of heterozygosity of a number of different mutations is an important driver of disease phenotype as well as therapy response (Kharazi et al., 2011). This is also demonstrated in our analysis of MPN patients, showing clear transcriptional differences between *JAK2* heterozygous and homozygous HSPCs in multiple patients with different disease phenotypes. Moreover, TARGET-seq has the advantage of combining scRNA-seq data and mutational analysis with index sorting, allowing tracing of cells back to canonical stem/progenitor cell hierarchies, revealing an aberrant HSPC phenotype in an MPN patient associated with presence of *JAK2* homozygous mutation. Furthermore, the reliable identification of wild-type cells made possible by TARGET-seq allows analysis of how tumors and treatment disrupt normal tissue residing cells. Such microenvironmental factors might underlie many aspects of tumor biology and therapy response. The high sensitivity mutation analysis made possible by TARGET-seq also allowed order of acquisition of mutations to be resolved, of importance as mutation order has been shown to correlate with disease phenotype and response to targeted inhibitors (Ortmann et al., 2015).

The throughput of this technique would typically allow the preparation of approximately 400 cells per week and of thousands of cells within a few months, in line with the numbers of cells analysed in published scRNA-seq datasets of tumors (Giustacchini et al., 2017; Tirosh et al., 2016a; Tirosh et al., 2016b). Whilst much higher throughput scRNA-seq techniques are available (Macosko et al., 2015; Zheng et al., 2017), these typically offer shallow coverage of only the 3’ or 5’ region of transcripts and lower molecular capture rates, and are consequently not designed for variant detection. The requirement of large numbers of cells would also prevent its use to study rare subpopulations of tumor cells such as CSCs. Consequently, there is a very important role for techniques that allow moderate throughput scRNA-seq (thousands of cells), generating libraries of high complexity with high-sensitivity mutation analysis, particularly when analyzing rare subpopulations of tumor cells. Moreover, analysis of alternative splicing patterns is made possible by the full-length scRNA-seq data generated by TARGET-seq, which is of paramount importance in most cancerous tissues (David and Manley, 2010) as well as many other diseases (Cooper et al., 2009), particularly as components of the spliceosome machinery are recurrently mutated in cancer (Kandoth et al., 2013).

In summary, TARGET-seq provides a powerful tool to resolve distinct molecular signatures of genetically distinct subclones of tumor cells and we anticipate this will pave the way for application of scRNA-sequencing for the definitive analysis of ITH and the identification and characterization of therapy resistant tumor subclones.

### Limitations

A potential limitation of TARGET-seq is that this approach does not support mutation discovery and relies on the analysis of known driver mutations, or mutations previously identified by other discovery-type methods. However, as the lysate is initially frozen and stored, this will routinely allow for mutational analysis from the same sample before subsequent processing of single-cells. Up to now, we have multiplexed primers to detect a total of twelve different mutations per single-cell. Whist this will be adequate to analyze key driver mutations in many tumors, for more genetically complex tumors, a more complex multiplexing strategy might be required. For very genetically complex tumors where potentially hundreds of different mutations need to be tracked, a whole genome/whole transcriptome approach may be more appropriate (Dey et al., 2015; Macaulay et al., 2015), albeit at the cost of reduced sensitivity of detection of those mutations (Hosokawa et al., 2017; Wang et al., 2014). Whilst TARGET-seq is currently optimized for full-length scRNA-seq analysis, this method could be easily and readily adapted to other single-cell RNA-sequencing technologies that are 3’ or 5’ biased such as SCRB-seq (Soumillon et al., 2014). Such adaptations could increase scalability and reduce sequencing costs associated with full-length techniques. In the current study, we have applied this technique to analyze hematopoietic tumors, however, this method could be broadly applied to the analysis of a range of cancers, and provides a powerful new tool to link transcriptional signatures with genetic tumor heterogeneity.

## Supporting information

## Acknowledgments

We thank the patients and staff in the National Cancer Research Network (NCRN), Dr. Deena Iskander and MDSBio study for samples, Dr. Nguyen Tran for laboratory management and banking of clinical samples and Dr. Neil Ashley for technical assistance. This work was funded by a Medical Research Council Senior Clinical Fellowship (MR/L006340/1) to A.J.M., a CRUK DPhil Prize Studentship (C5255/A20936) to A.R-M. and the MRC Molecular Haematology Unit core award (A.J.M. and S.E.W.J.; MC_UU_12009/5). This work was also supported by Neil Ashley in the WIMM Single Cell Facility and the MRC-funded Oxford Consortium for Single-cell Biology (MR/M00919X/1) and the Oxford NIHR Biomedical Centre based at Oxford University Hospitals NHS Trust and University of Oxford. The views expressed are those of the author(s) and not necessarily those of the NHS, the NIHR, the Department of Health or the NIH. The authors acknowledge the contributions of the WIMM Flow Cytometry Facility, supported by the MRC HIU; MRC MHU (MC_UU_12009); NIHR Oxford BRC and John Fell Fund (131/030 and 101/517), the EPA fund (CF182 and CF170) and by the WIMM Strategic Alliance awards G0902418 and MC_UU_12025.

## Author Contributions

A.R-M. designed, performed and analyzed experiments, performed bioinformatic analyses and contributed to writing the manuscript. G.B. developed method automation protocols. G.B., S.A.C, B.J.P, V.A.D., E.L. and N.S. assisted with experiments. B.J.P. analysed data. E.L., B.P. and N.S. processed clinical samples. E.L., B.P, N.S. and A.H. provided clinical information. S.M and N.B. provided bioinformatic pipelines. A.G. provided protocols and technical input. S.E.W.J. provided input in experimental design and analysis, and provided input on writing the manuscript. S.T. designed and supervised bioinformatic analyses. A.J.M conceived and supervised the project, designed and analyzed experiments and wrote the manuscript. All authors read and approved the submitted manuscript.

## STAR Methods

### CONTACT FOR REAGENT AND RESOURCE SHARING

Further information and requests for resources and reagents should be directed to and will be fulfilled by Adam Mead.

### Cell lines

K562, MOLT4 and JURKAT cells were obtained from the American Type Culture Collection (ATCC). NALM6 cells were obtained from the German Collection of Microorganisms and Cell Cultures (DSMZ). SET2 cells were kindly provided by Dr. Jacqueline Boultwood and Dr. Andrea Pellagatti (Radcliffe Department of Medicine, University of Oxford). All cell lines were maintained in culture in RPMI-1640 supplemented with 10 % Fetal Calf Serum (FCS) and antibiotics. Cell lines were authenticated by targeted sequencing of known mutations.

### Banking and processing of human samples

Patients and normal donors provided written informed consent in accordance with the Declaration of Helsinki for sample collection and use in research under the INForMeD Study (REC:199833, University of Oxford). Cryopreserved peripheral blood and bone marrow mononuclear cells were thawed and processed for flow cytometry analysis as previously described (Woll et al., 2014). A summary of patients and normal donors samples used for analysis can be found in Table S3.

### Fluorescent activated cell sorting (FACS) staining and single-cell isolation

Single cell FACS-sorting was performed as previously described (Giustacchini et al., 2017), using BD Aria III or BD Fusion I instruments (Becton Dickinson). All the experiments involving isolation of human hematopoietic stem and progenitor cells (HSPCs) included single color stained controls (CompBeads, BD Biosciences) and Fluorescence Minus One controls (FMOs). Lineage-CD34+ cells were sorted and indexed for CD38, CD90, CD45RA and CD123 markers, which allowed us to record the fluorescence levels of each marker for each single cell. The full list of antibodies used for HSPCs immunophenotyping and isolation can be found in Key Resources; 7-aminoactinomycin D (7-AAD) was used for dead cell exclusion. Briefly, single cells directly sorted into 96-well PCR plates containing 4-4.2 µL of lysis buffer. K562 cells were sorted into the lysis buffer described in Table S1a. JURKAT, MOLT4, NALM6, SET2 and HSPC were sorted into lysis buffers described in Table S1b. Flow cytometry profiles of the HSPC compartment of normal donors and patient samples (Figure S4) were analyzed using FlowJo software (version 10.1).

### cDNA synthesis

For K562 cells, RT and PCR steps were performed as described in Table S1a, using 18 cycles of PCR amplification. For JURKAT, MOLT4, NALM6, SET2 cells and HSPCs, RT and PCR steps were performed as described in Table S1b, using 20 cycles of PCR amplification for cell lines and 22 cycles of amplification for HSPCs. ERCC Spike-in (Ambion, #4456740) was used at a final concentration of 1:40,000,000 in 10 µL of RT mix for cell lines (K562, MOLT4, JURKAT, SET2, NALM6) and 1: 200,000,000 for HSPCs. The sequences of the primers used in the RT and PCR steps, for whole transcriptome and targeted retrotranscription and cDNA amplification, can be found in Table S2a. Primers were designed to amplify amplicons 250-700 bp long and specificity was checked against RefSeq and human genome assembly databases using PrimerBlast. mRNA and cDNA primers were designed to amplify coding regions whereas gDNA primers were designed to bind at least to one intronic region. After PCR, 15 µL from a total of 25 µL cDNA mix were diluted with 11 µL of water and purified using 16 µL of Ampure XP Beads (0.6:1 beads to cDNA ratio; Beckman Coulter), and resuspended in a final volume of 8 µL of EB buffer (Qiagen). The quality of cDNA traces was checked using a High Sensitivity DNA Kit in a Bioanalyzer instrument (Agilent Technologies). The remaining 10 µL of the cDNA mix were used for single-cell genotyping or stored at −20 C.

### Targeted NGS single-cell genotyping

After cDNA synthesis, 1 µL aliquot from each single cell derived library was used as input to generate a targeted and Illumina-compatible library for single cell genotyping. The preparation of single cell genotyping libraries involves 2 PCR steps (See Detailed Protocol). In the first PCR step, target specific primers (Table S2b) attached to Access Array adaptors (Fluidigm; Forward adaptor: ACACTGACGACATGGTTCTACA; Reverse adaptor: TACGGTAGCAGAGACTTGGTCT) are used to amplify the target regions of interest. Barcoding primers were designed to specifically amplify cDNA or gDNA, amplifying annotated coding regions in the case of cDNA amplicons and at least one intronic region in the case of genomic DNA amplicons. In the second PCR step (See Detailed Protocol), Illumina compatible adaptors containing single-direction indexes (Access Array™ Barcode Library for Illumina® Sequencers-384, Single Direction, Fluidigm) are attached to pre-amplified amplicons from the first PCR to generate single-cell barcoded libraries. All NGS single-cell genotyping library preparation steps are automated in a Biomek FxP Workstation. Amplicons were pooled, purified with Ampure XP beads (0.8:1 ratio beads to product), quantified using Quant-iT Picogreen (Thermo Fisher Scientific) and pools were diluted to a final concentration of 4 nM.

Up to 384 single cells were sequenced on a MiSeq (Illumina) instrument, with 150 bp paired-end reads. Reads were aligned to GRCh37/hg19 using STAR with default settings (version 2.4.2a) and cDNA/gDNA amplicons were separated into different bam files using a custom Perl script, extracting reads matching the different primer sequences used for targeted PCR barcoding. This allowed us to obtain independent mutational information from cDNA and gDNA. Variant calling was performed using mpileup (samtools version 1.1, options --minBQ 30, --count-orphans, --ignore overlaps) and results were summarized with a custom Python script. Thresholds for the detection of each amplicon were set based on non-template controls and thresholds for mutation calling were based on WT controls (1-5 % of the reads, Figures S2b, S2c). Both non-template and WT controls were routinely processed in parallel to test samples. Importantly, none of the tested mutations were detected in any control cells (n=112) in any of the experiments, implying that the false positive rate of variant calling with the cut-offs used is effectively zero. For experiments involving isolation of HSPCs, single cells where one of the targeted amplified genes tested failed to be detected by both gDNA and mRNA were excluded from analysis.

### Nextera XT library preparation and Illumina whole transcriptome sequencing

Bead-purified cDNA libraries were used for tagmentation with Nextera XT DNA Kit (Illumina) using one fourth of the original volume as previously described (Giustacchini et al., 2017). 4nM libraries were sequenced on a NextSeq instrument with 75 bp single-end reads.

### Single cell RNA-sequencing data pre-processing

RNA-sequencing reads were trimmed for Nextera adaptors with TrimGalore (version 0.4.1) and aligned to the human genome (hg19) using STAR with default settings (version 2.4.2a). RefSeq gene model was used as the reference for gene expression quantification. Counts for each RefSeq gene were obtained with FeatureCounts (version 1.4.5-p1; options --primary) and were normalized to reads per kilobase per million mapped reads (RPKM). Genes with RPKM values less or equal than 1 were considered non-detected and expression values for these genes were converted to zero. We further normalized RPKM expression values into the log2 scale. QC filtering was performed using the following parameters: percentage of reads mapping in exons > 50 %, percentage of mapped reads > 50 % and number of detected genes per cell (RPKM>=1) > 6000 for JURKAT and SET2 cells, > 5000 for K562 cells and > 1500 for primary HSPCs. For cell lines, we excluded 8 cells after applying these QC filters (5.3 %) and for HSPCs, 33 cells (6.1 %).

### Whole transcriptome variant calling from single cells

Bam files from 38 single HSPC (Figure 1e) or 48 single K562 cells (Figure 1c) were merged using samtools to computationally create a single cell ensemble. LoFreq software (Wilm et al., 2012) was used for variant calling in the single cell ensemble. Heterozygous regions across the transcriptome (AF> 0.05 of the minor allele, Allele Frequency) were used for variant calling in each individual cell, requiring a minimum coverage of 10 reads and minimum base quality of 30. A SNV was considered heterozygous if AF> 0.05 and homozygous if AF<0.05.

### Mutational analysis from RNA-sequencing reads

Variant calling from raw RNA-sequencing reads was performed using mpileup (samtools version 1.1, options --minBQ 30, --count-orphans, --ignore overlaps) and results were summarized with a custom Python script. Thresholds for the detection of amplicons were set at 30 reads per position (Figure S2c), in line with variant calling guidelines (Sims et al., 2014).

### Dropout frequency and library bias calculation

The frequency of dropout for a given gene was calculated as the percentage of cells from a specific condition (SMART-seq2 or SMART-seq+) in which the gene is not detected (RPKM<1), as compared to the average expression of that gene in K562 bulk samples (6 replicates of 100 cells each; 3 replicates per chemistry). Library bias was calculated as the ratio between the mean rpkm of the top 10 % expressed genes in the library and the mean rpkm of all genes.

### Transcript coverage

Normalized transcript coverage was calculated using “geneBody_coverage.py” script from RSeQC package (http://rseqc.sourceforge.net), using 4040 housekeeping genes.

### Cell cycle phase assignment and correction

Cell cycle phase for HSPCs from patient IF0602 was performed using “cyclone” function implemented in the scran package in R (Scialdone et al., 2015), using a geneset of cell-cycle associated human genes. Cell cycle effect was removed using limma package in R, using cell cycle phase identified by “cyclone” function.

### Single cell clustering and dimensionality reduction

Principal component analysis (PCA) was performed using log2-transformed RPKM values of 13793 expressed genes (limit of detection RPKM>=1) with “prcomp” function in R (Figure S6a). T-distributed stochastic neighbor embedding (tSNE) was performed for single cells from patient IF0602 using ‘Rtsne’ package, the implementation of the method in R, with “perplexity=20”. We used log2-transformed RPKM values of 13793 expressed genes with a limit of detection of RPKM>=1 (Figure S6b) or 5129 highly variable genes (Figure S6d), identified as previously described (Giustacchini et al., 2017). Unsupervised hierarchical clustering of Spearman’s correlations using 5129 highly variable genes was performed using “cor” function in R (Figure S6c). Cell cycle labelled (Figure S6e) or cell cycle corrected (Figure S6f) tSNE were performed using the same 5129 highly variable genes as in Figure S6d, with “perplexity=20”. SC3 software (Kiselev et al., 2017) was also used to analyze the subclonal composition of patient IF0602, using default parameters and k=2 (as there are two groups of genetically distinct cells; Supplementary Figure 6g). Unsupervised hierarchical clustering of the top 500 differentially expressed genes between *JAK2* WT and *JAK2V617F* mutant cells from patient IF0602 was performed using ‘pheatmap’ package in R (Figure S6h).

### Random forest analysis

Random forest analysis was performed using ‘randomForest’ package in R (ntree=2000), trained on the genotypes of single cells. Only genotypes with more than five cells were included in this analysis. Expression matrix was batch and donor-corrected using ‘limma’ package in R. The top 2000 genes identified by the random forest analysis (MeanDecreaseGini > 0.041) were used for the tSNE representation in Figure 5c (perplexity=20). Clustering of cells was stable when selecting from 500 to 5000 top genes from the random forest analysis.

### Gene-set enrichment analysis

GSEA was performed using GSEA software (http://software.broadinstitute.org/gsea) with default parameters and 1000 permutations on the phenotype. Gene sets used for the analysis were downloaded from MSigDB or relevant studies (Table S4). Single Sample GSEA (ssGSEA) was performed using ssGSEA Projection Module (https://genepattern.broadinstitute.org) with default setttings and combine mode ‘combine.off’. A projection of ssGSEA results is shown in Figure 5b.

### Differential expression analysis

Differentially expressed genes were identified using a combination of non-parametric Wilcoxon test and Fisher’s exact test as previously described (Giustacchini et al., 2017) (Table S5). Significant genes were selected on the basis of adjusted *P* value < 0.1 and absolute log2(fold-change)>0.5. P-values corresponding to Wilcoxon test and Fisher’s exact test are shown on the top or each bar in Figures 4c and 4e. Beeswarm plots from selected genes were generated using “beeswarm” package in R.

### Statistical analysis

Unpaired Student t-test with Welch’s correction was used for the comparisons in Figure 1a, Figure 1b, Figure S1a, Figure S1b and Figure S3a.

### Code availability

R, Perl and Python scripts used for the analysis would be available upon publication.

### Data availability

Single cell RNA-sequencing data would be available at the NCBI’s GEO data repository upon publication. Single cell targeted sequencing data is available at the NCBI’s SRA data repository upon publication.

---

**Figure S1.**
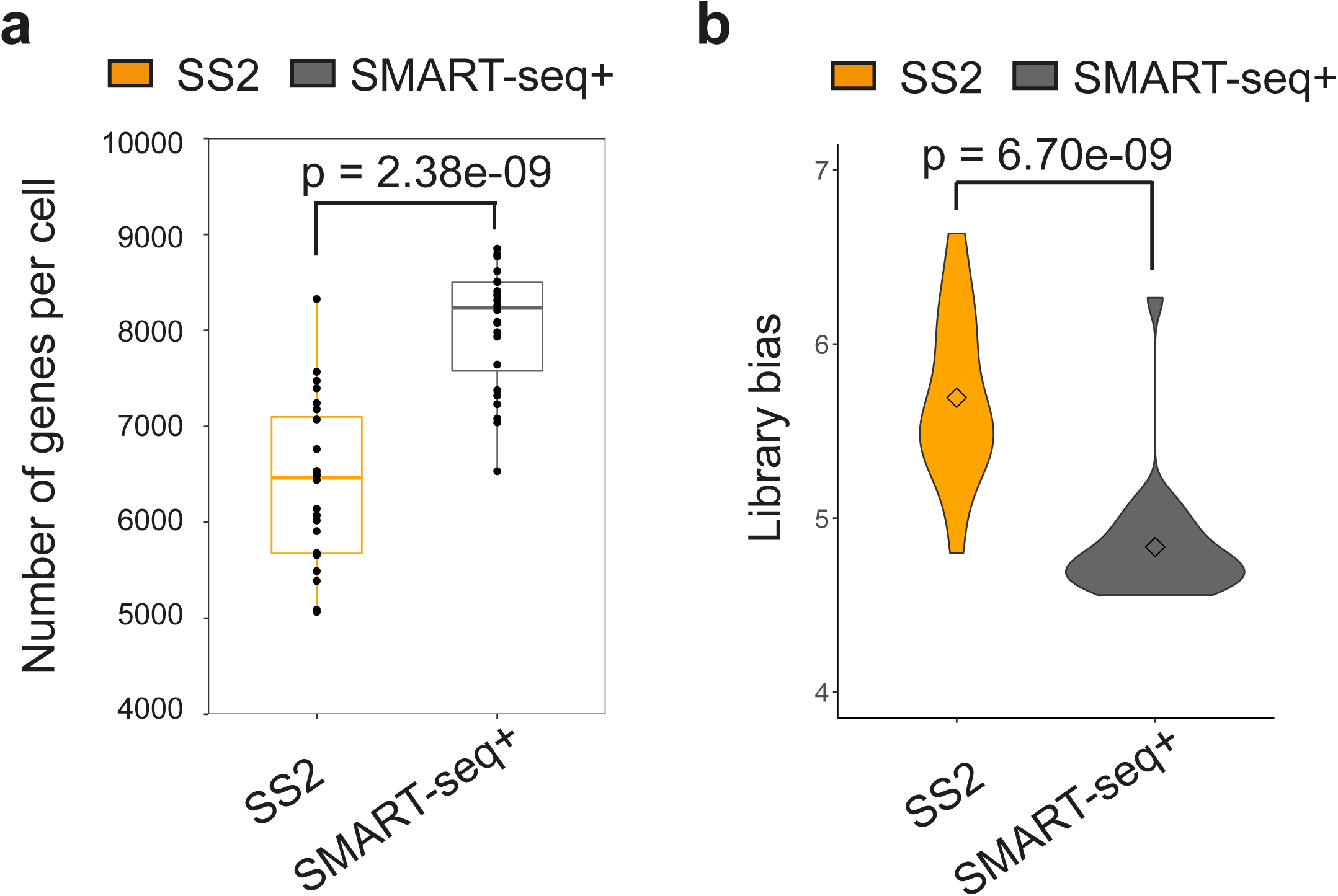
related to Figure 1. SMART-seq+ is an optimized single-cell RNA-sequencing chemistry. **(a)** Number of detected genes per cell in K562 cells processed using Smart-seq2 or optimized SMART-seq+ chemistry (n=48, 24 cells per chemistry from three independent experiments). P-value from two-tailed unpaired Student t test is shown on the top of the graph. **(b)** Library bias per chemistry, calculated as the ratio between the mean rpkm values of the top 10% expressed genes and the mean rpkm for all genes expressed in the library, using the same 48 cells as in (a). P-value from two-tailed unpaired Student t test is shown on the top of the graph. Points represent the mean for each group.

**Figure S2.**
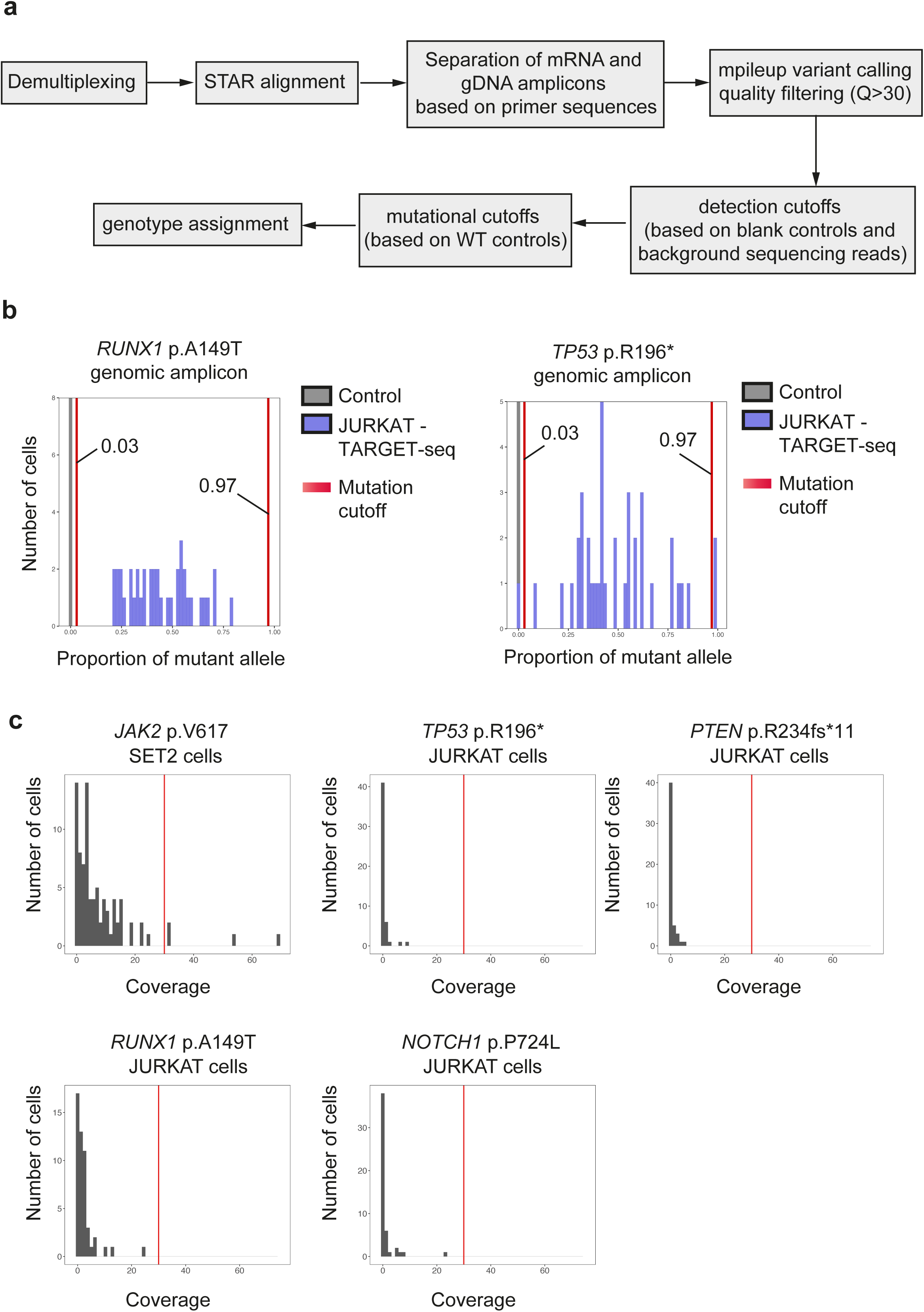
related to Figure 2. Targeted pre-amplification and sequencing of mRNA and gDNA amplicons dramatically increases the sensitivity of mutation detection. **(a)** Schematic representation of the pipeline used for variant calling of targeted next generation sequencing from single cells. **(b)** Representative examples of allelic distributions and mutational cut-offs of *RUNX1* p.A149T and *TP53* p.R196* mutations in JURKAT cells. Red lines represent mutation cut-offs (<3% and >97% of reads mapping to mutant allele). **(c)** RNA-sequencing coverage of *JAK2* mutation in SET2 cells and *TP53, NOTCH1, RUNX1* and *PTEN* mutations in JURKAT cells. The y-axis represents the number of cells against their coverage for each mutation in the x-axis. Red line represents a coverage threshold of 30.

**Figure S3.**
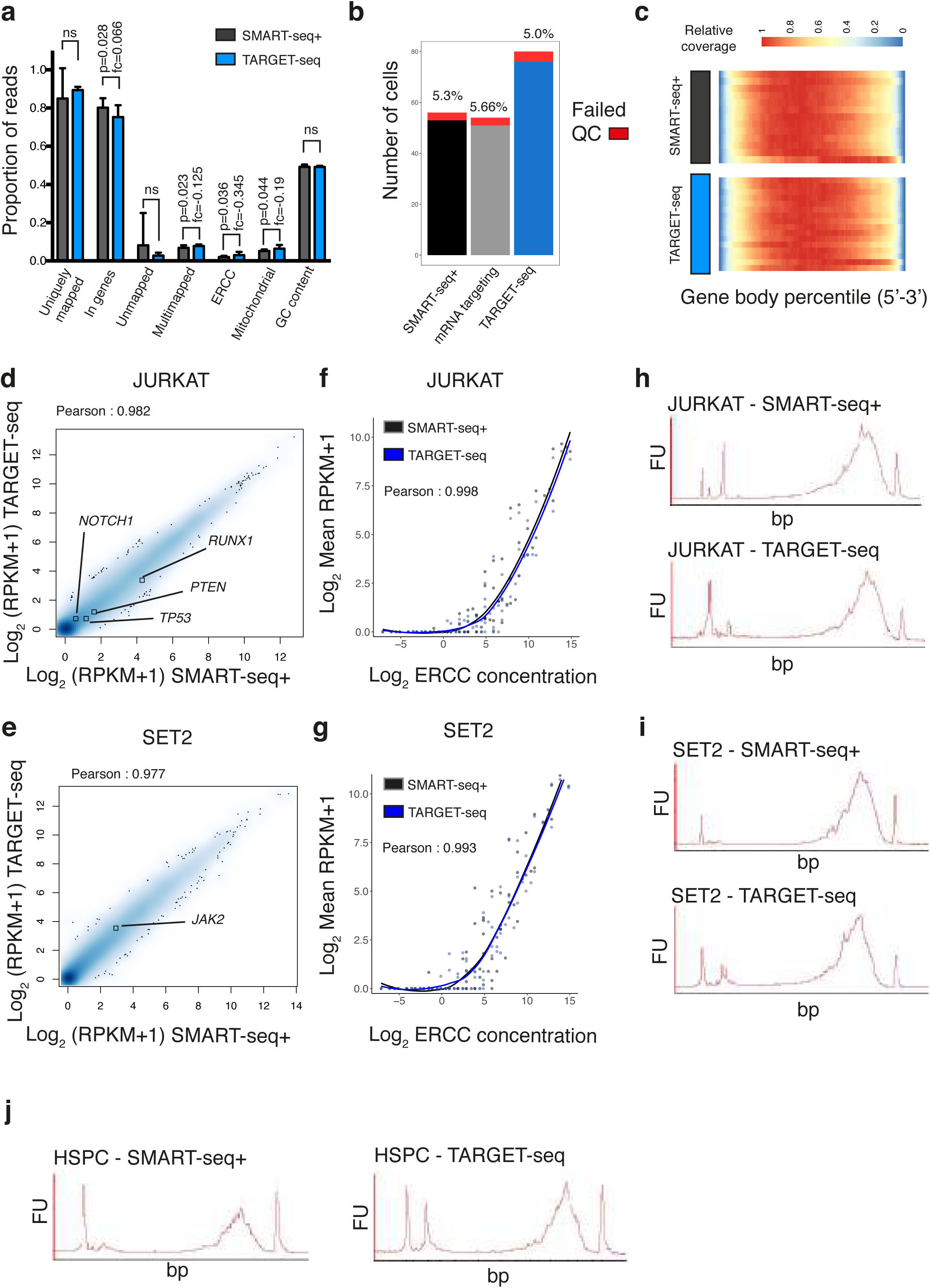
related to Figure 3. TARGET-seq produces unbiased whole transcriptome data from single cells. **(a)** Sequencing statistics of single cell libraries from HSPCs processed using SMART-seq+ or TARGET-seq. The bar graph represents the proportion of reads for each sequencing statistic and condition, and error bars represent standard deviation of the mean. P-values from two-tailed Student t test and fold change values for each sequencing statistic are shown on the top of each pair of bars. **(b)** Number of cells passing or failing QC per method. The percentage of cells failing QC for each method is shown on the top of each bar. **(c)** Normalized transcript coverage from single HSPCs processed using SMART-seq+ or TARGET-seq methods, using a list of 4040 housekeeping genes. **(d,e)** Whole transcriptome Pearson’s correlation between SMART-seq+ and TARGET-seq ensembles (mean RPKM values per condition) in JURKAT **(d)** and SET2 cells **(e)**. The expression values for the genes targeted are highlighted in each cell type. **(f,g)** Pearson’s correlation between mean ERCC expression values from SMART-seq+ and TARGET-seq in JURKAT cells **(f)** and SET2 cells **(g)**. **(h-j)** Bioanalyzer traces of representative cDNA libraries synthetized using SMART-seq+ or TARGET-seq in JURKAT **(h)**, SET2 **(i)** or HSPCs **(j)**.

**Figure S4.**
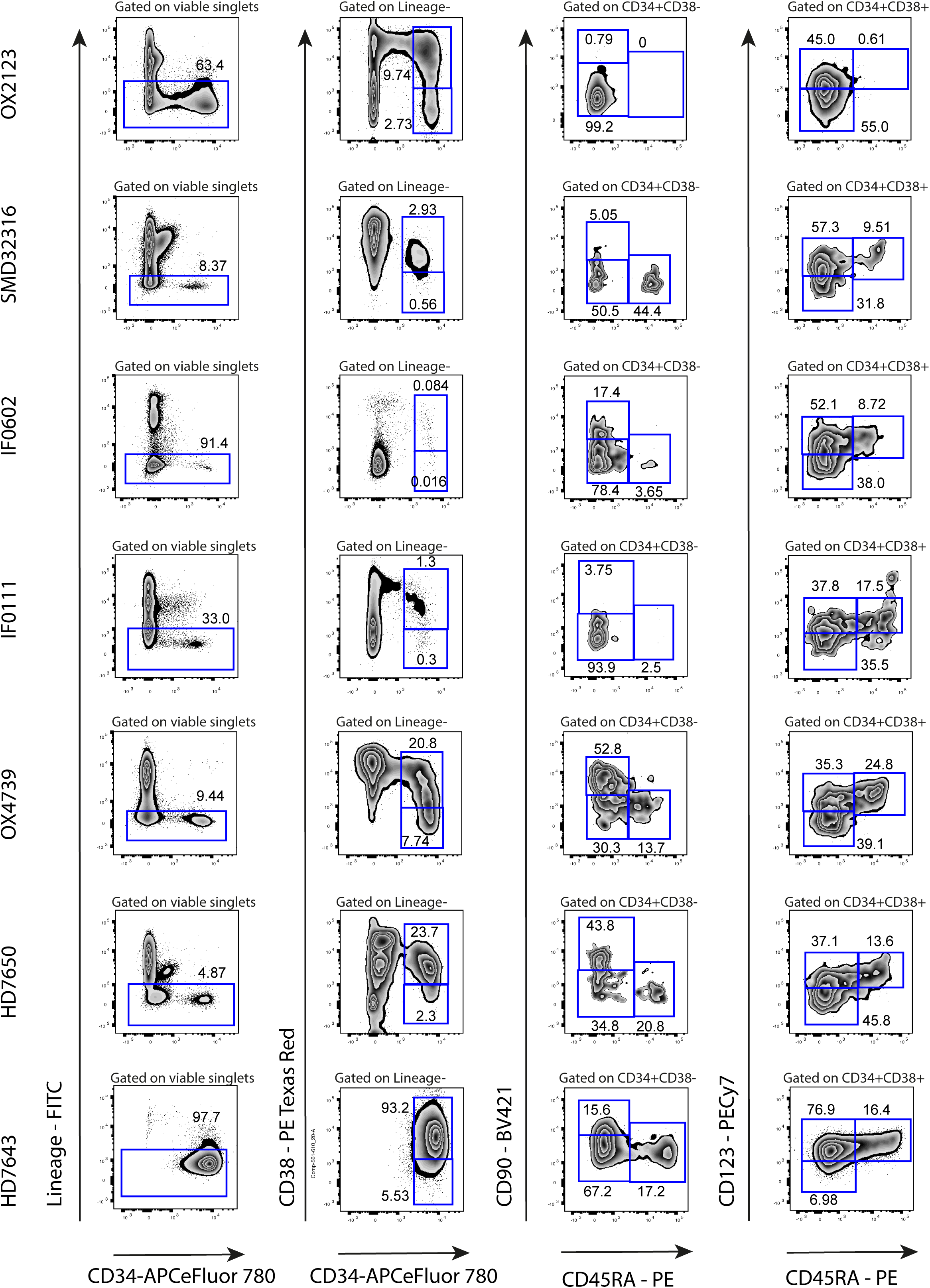
related to Figures 4 and 5. Representative flow cytometry profiles and gating strategies for all five patients analyzed and two normal donors. Numbers represent percentage of gated cells. Antibodies used for HSPC isolation are listed in Key Resources Table.

**Figure S5.**
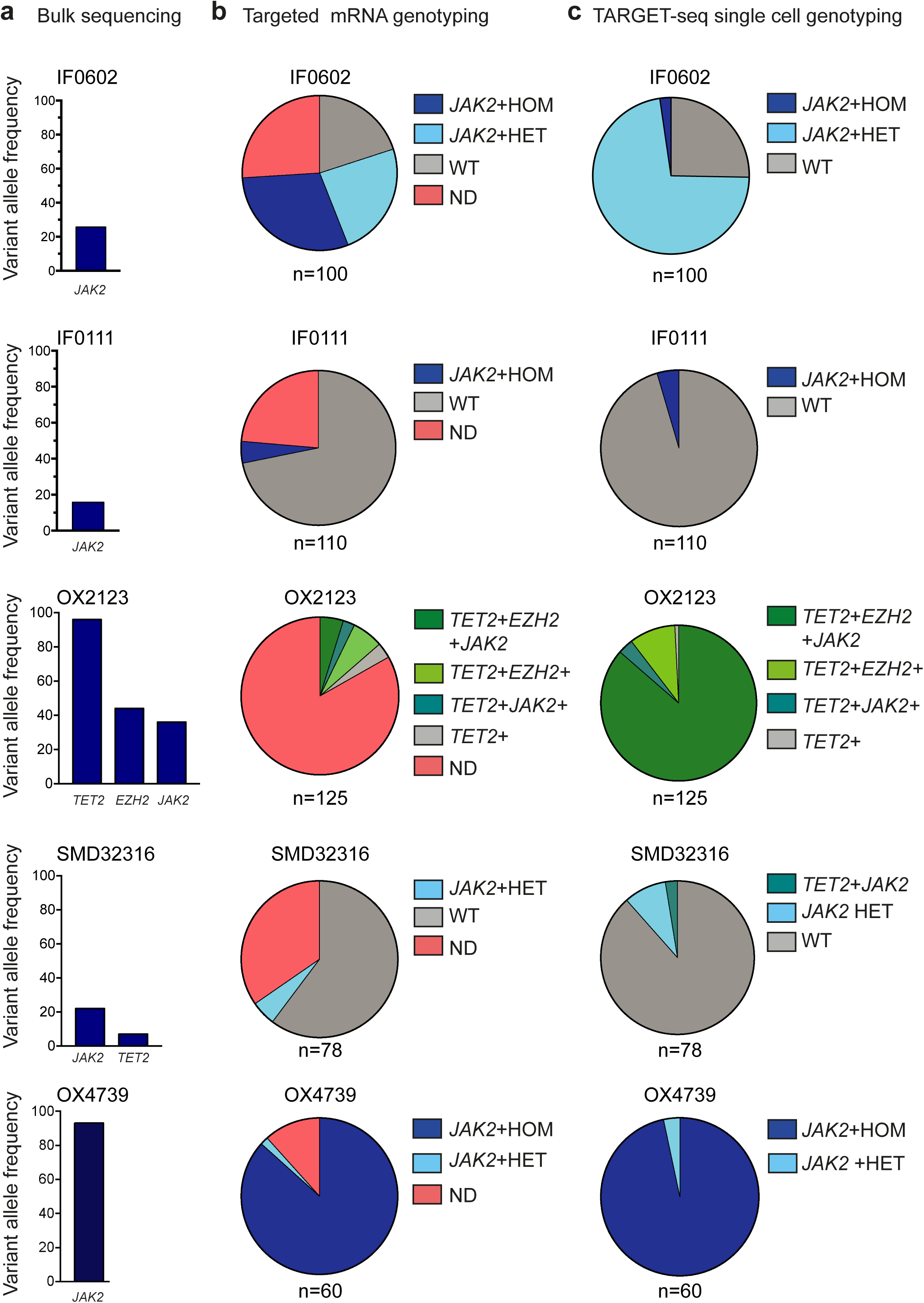
related to Figures 4 and 5. Single cell sequencing reveals subclonal structures in the HSPC compartment that could not be inferred through bulk sequencing. **(a)** Variant allele frequency of mutations identified in each patient by bulk sequencing of total MNCs using a clinically validated myeloid panel. Patient’s code (Table S3) indicates mutations identified in each patient. **(b,c)** Subclonal composition of each patient’s HSPC compartment identified by single cell genotyping using mutational information from targeted mRNA amplicons (mRNA targeting) or (c) TARGET-seq. Total number of cells analysed is shown under the pie chart for each patient, and each patient is labeled according to the code provided in Table S3. ND (Not Detected) represents cells in which at least one of the amplicons was not detected. Only cells passing genotyping QC for each patient are show.

**Figure S6.**
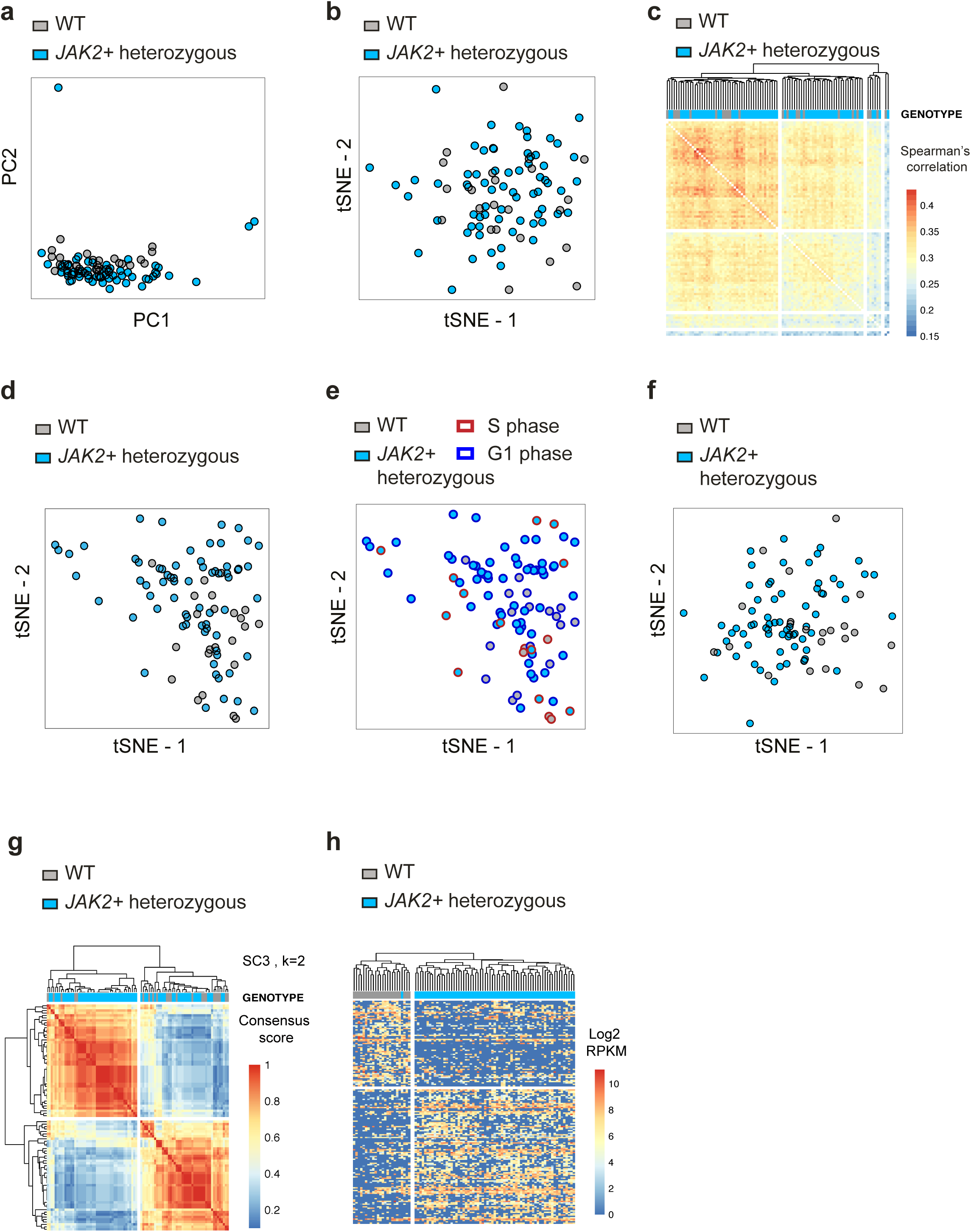
related to Figure 4. Computational methods commonly used for single cell RNA-sequencing analysis are unable to distinguish between genetically distinct JAK2V617F mutant subclones. **(a,b)** PCA **(a)** or tSNE **(b)** analysis of JAK2V617F heterozygous mutant and WT cells from patient IF0602 using 13793 expressed genes at RPKM>=1. Cells are colored according to genotype. Hierarchical clustering of Spearman’s correlations between single cells from patient IF0602 using 5129 highly variable genes. **(d,e)** tSNE analysis of patient IF0602 using 5129 highly variable genes, coloring cells according to genotype **(d)** or cell cycle stage **(e)**, **(f)** tSNE analysis of cells from patient IF0602 removing cell cycle phase effect using limma. **(g)** SC3 k-means clustering of patient IF0602 using k=2, as this patient has two genetically distinct subclones. **(h)** Hierarchical clustering of single cells from patient IF0602 using the top 500 differentially expressed genes between *JAK2* WT and *JAK2V617F* mutant cells.

## SUPPLEMENTARY TABLES

**Table S1, related to Figure 1, Figure S1, Figure 2 and Figure 3.** Reagents and conditions used for cDNA synthesis of SMART-seq2, SMART-seq+, mRNA targeting and TARGET-seq libraries.

**Table S2, related to Figure 2.** Primers used for amplification of mRNA and gDNA amplicons during RT, PCR and barcoding PCR steps.

**Table S3, related to Figures 4 and 5, Figures S4 and S5.** Summary of patient samples (n=5) and normal donors (n=2) used for the study.

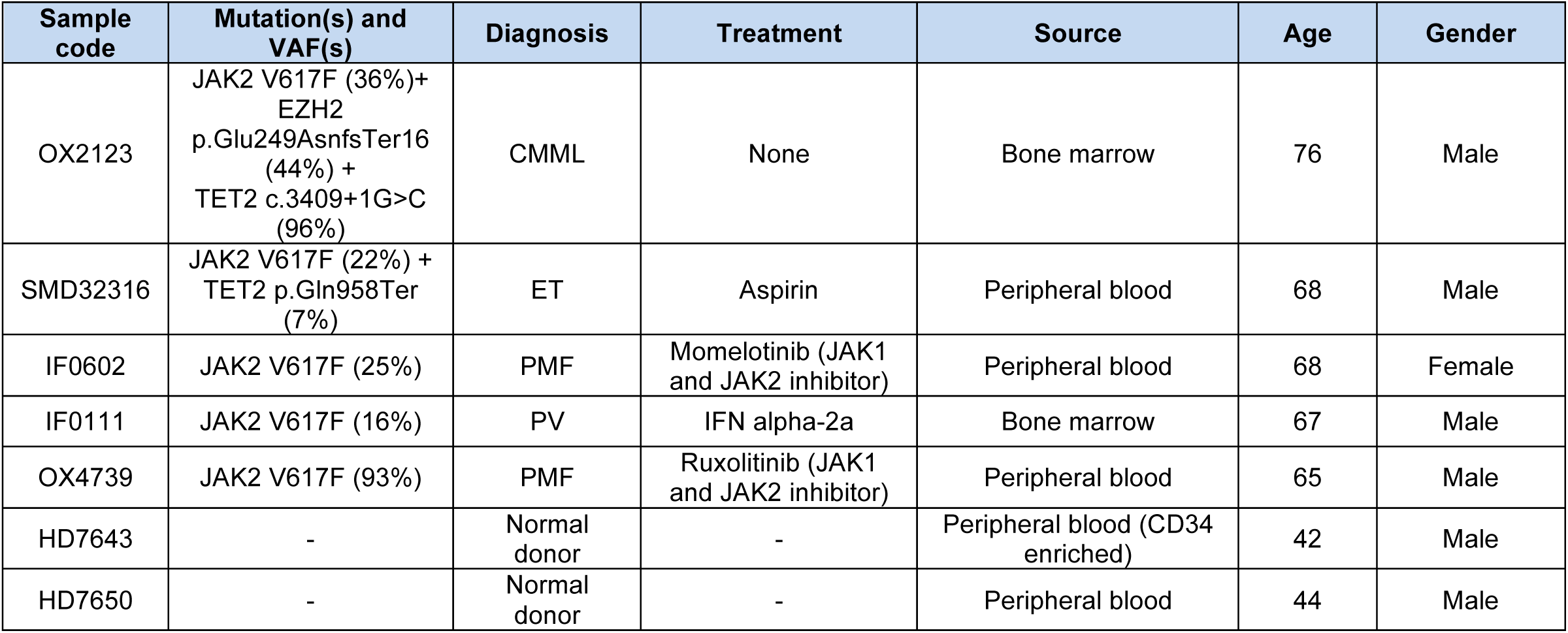

**Table S4, related to Figure 4 and Figure 5**. Genesets used for GSEA and ssGSEA analysis.

**Table S5, related to Figure 4.** Differentially expressed genes and GSEA analysis of *JAK2V617F* mutant, *JAK2* WT cells and cells from normal donors.

**Table S6, related to Figure 5.** GSEA analysis of *EZH2, TET2* and *JAK2V617F* mutant cells compared to cells from normal donors.

