## Supplementary material for "Unraveling intratumoral heterogeneity through high-sensitivity single-cell mutational analysis and parallel RNA-sequencing"

### FULL LENGTH TARGET-SEQ DETAILED PROTOCOL – 96 well plates

#### Materials

- 96-well PCR plates (AB-0900)
- 384 well PCR plates (FrameStar, 4titude, 4ti-0384/C)
- Corning® 96 Well TC-Treated Microplates (Cat. No. CLS3595-50EA)
- RNase Free Microfuge Tubes (Invitrogen, AM12400)
- V-shaped 96 well plate (Cat. No. P-96-450V-C, Axygen)
- PCR film (MicroAMp Clear Adhesive Film, Cat. No. 4306311)
- Aluminium Sealing Film (StarLab, E2796-0792)
- 96-well magnetic stand
- Protease (Qiagen #19155)
- Triton X-100 (Sigma-Aldrich # T8787)
- RNase inhibitor (TAKARA #2313A)
- dNTPs (Life Technologies #R0192)
- UltraPure DNAse/RNAse-Free Distilled Water (Life Technologies, #10977035)
- EB Buffer (Qiagen Cat No./ID: 19086)
- Ethanol
- ERCC aliquot (Ambion #4456740)
- SMARTScribe enzyme (Clontech - Cat. No. 639537).
- SeqAMP enzyme (638509 – Clontech).
- RT-PCR Grade Water (Life Technologies - AM9935).
- Ampure XP beads (Beckman Coulter; Cat. No. A63881)
- Pre-amplification primers: oligodT-ISPCR primer, mRNA target-specific primers, TSO-LNA (RNase free HPLC purification); gDNA target-specific primers, cDNA target-specific primers and ISPCR primers (HPLC purification).
- PCR1 primers: CS1/CS2-target specific primers for gDNA and cDNA PCR1 barcoding (Cartridge purification).
- Sequencing primers: CS1, CS2, CS1rc, CS2rc (custom LNA primers from Exiqon-Qiagen; HPLC purification).
- Nextera XT Kit Library Preparation Kit (Cat. No.15032354, Illumina) including i7 indexes and i5 indexes (Nextera XT Index Kit, FC-131-1001, Illumina)
- KAPA 2G Robust HS PCR Kit (#KK5517)
- FastStart High Fidelity PCR System (Roche, REF:04738292001)
- Access Array™ Barcode Library for Illumina® Sequencers-384, Single Direction (Fluidigm, Cat. No. 100-4876).

#### Sorting and lysis – Timing: variable

1. First, prepare sufficient lysis buffer for the required number of cells for each experiment, plus 10% dead volume. Aliquot the lysis buffer (containing oligodT-ISPCR primer) into each well of a 96-well PCR plate (Thermo Scientific #AB-0900) in a clean environment dedicated to ‘pre-amplification’ work only. Cover with a PCR film and keep on ice/in the fridge until use. Lysis buffer should be prepared fresh on the day of sorting, maximum a few hours before use.

2. Prepare the sorter for single-cell sorting. Use single cell purity mode and keep the event rate low (less than 1000/s).

| **Lysis** | **1 cell** | **Storage** | **Cat. No./Supplier** |
|---|---|---|---|
| Triton 0.4% | 1.9 µL | -20 °C | Sigma-Aldrich # T8787, resuspended in water |
| RNAse Inhibitor | 0.1 µL | -20 °C | TAKARA #2313A |
| dNTPs (10 mM) | 1 µL | -20 °C | Life Technologies #R0192 |
| Oligo-dT-ISPCR (10 uM) | 1 µL | -20 °C | Biomers (custom oligo, HPLC purified) |
| Protease (1.09 AU/mL in water) | 0.1 µL | +4 °C | Qiagen #19155; resuspend in UltraPure DNAse/RNAse-Free Distilled Water (Life Tech, #10977035) |
| ERCC RNA spike-in mix (1:2e6) | 0.1 µL | -80 °C(single use aliquot) | Ambion #4456740 |
| **TOTAL** | 4.2 µL | | |

3. Check sorter alignment: use a 96 well tissue culture flat-bottom plate (Corning) and sort one fluorescent bead per well. Check under a fluorescent microscope that there is only one bead per well and that the position of the bead is centered. Note: don’t add any media into the plate so that the bead stays in the place it was deposited by the sorter.

4. Use a 96-well PCR plate (Thermo Scientific #AB-0900, same model as the one in which cells will be sorted) covered with a PCR film, and sort 50 cells in positions 1A, 1H, 12A and 12H. Droplets should be positioned in the centre of the wells if the sorter is correctly aligned; if not, make necessary adjustments until drops are falling perfectly into each well.

5. After this initial check, remove the PCR film from the 96 PCR plate and sort 50 cells into columns 1 and 12 (wells 1A, 1B, 1C, 1D, 1E, 1F, 1G, 1H and 12A, 12B, 12C, 12D, 12E, 12F, 12G, 12H): drops should now be deposited at the very bottom of each well with no traces of liquid been left in the sides of each well. If correct, alignment checks are now complete.

6. Perform a purity sort of desired populations.

7. Sort cells directly into a 96 well PCR plate (Thermo Scientific #AB-0900) containing the lysis buffer, cover the plate with an aluminium PCR film (StarLab), spin down the plate and incubate 5 minutes at room temperature to allow for protease digestion. If sorting time is longer than 10 minutes, there is no need to incubate the plate further.

8. Put the plate directly into dry ice and store at -80 °C up to 1-2 months. (Processing plates after 3 months of -80 °C storage has shown decreased yield and/or signs of RNA degradation).

**Heat inactivation, cDNA synthesis and amplification (RTPCR) – Timing: 6.5 hours; 1.5 hours hands-on time**

9. Transport the plate(s) and TSO-LNA aliquot from -80 C storage on dry ice to a 'pre-amplification' dedicated workspace/clean room.

10. Thaw the 5X Buffer, RT-PCR Water and any mRNA targeting primers that you might add to the mix. These can be thawed at room temperature. Aliquot them into an RNase free tube to prepare a master mix for the RT step as per the table below. RNase inhibitor, TSO-LNA and SMARTScribe enzyme will be added to the mix during heat inactivation step.

| RT | 1 cell (µL) | Storage | Cat. No. |
|---|---|---|---|
| **Buffer 5X** | 2.00 µL | -20 °C | Clontech - Cat. No. 639537 (delivered with enzyme) |
| **RNase Inhibitor** (wait until the 72C step to add it) | 0.25 µL | -20 °C | TaKara - 2313A |
| **TSO-LNA (100 µM)** (wait until 72C step to add it) | 0.10 µL | -80 °C (aliquot into single use aliquots to avoid freeze/thaw cycles) | Custom TSO-LNA oligo from Exiqon-Qiagen (same as Picelli et al., 2013) |
| **RT-PCR Grade Water** | Variable | -20 °C | Life Tech - AM9935 |
| **mRNA primers** (0.035 µL of each primer from a 200 uM stock) | Variable | -20 °C | Custom HPLC purified primers from biomers.net; resuspend in RNAse Free TE/water |
| **SMARTScribe** (wait until 72C step to add it) | 1.00 µL | -20 °C | Clontech - Cat. No. 639537 |
| **TOTAL** | 5.60 µL | | |
| **TOTAL (cumulative)** | 9.80 µL | | |

11. Thaw the sample plate on ice/4°C and incubate 15 minutes at 72 °C. This step will inactivate the protease included in the lysis buffer so it doesn't interfere with any subsequent enzymatic steps.

12. During the heat inactivation time, add the RNAse Inhibitor, TSO-LNA and RT enzyme to the master mix on ice/cold block. Vortex and spin down.

13. After heat inactivation, take the plate out of the thermocycler, spin down and place into ice/cold rack. Aliquot 5.6 µL of RT mix into each well and carefully seal the plate with a PCR film (MicroAMp Clear Adhesive Film, Cat. No. 4306311). Note: t is essential that this step is performed within 5-7 minutes to avoid RNA degradation.

14. Spin down and run the following program in the thermocycler:

| Temperature | Time | Cycles |
|---|---|---|
| 42 C | 90 min | 1 |
| 50 C | 2 min | 10 cycles |
| 42 C | 2 min | |
| 70 C | 15 min | 1 |
| 4 C | HOLD | - |

15. Fifteen minutes before the RT program finishes, start thawing reagents to prepare the PCR master mix.

| PCR | 1 cell (µL) | Storage | Cat. No. |
|---|---|---|---|
| **2X Buffer** | 12.50 µL | -20 °C | Life Tech - AM9935 (delivered with enzyme |
| **ISPCR (10 µM)** | 0.125 µL | -20 °C | Custom HPLC oligo from biomers.net (same as Picelli et al., 2013) |
| **RT-PCR Water** | Variable | -20 °C | Life Tech - AM9935 |
| **SeqAMP Enzyme -** wait until RT is about to finish to add | 0.50 µL | -20 °C | 638509 - Clontech |
| **cDNA primers -** (0.035 µL from each primer from 20 µM stock) | Variable | -20 °C | Custom HPLC purified primers from biomers.net; resuspend in RNAse Free TE/water |
| **Genomic primers** (0.1 µL from each primer from a 200 µM stock) | Variable | -20 °C | |
| **TOTAL** | 15.00 µL | | |
| **TOTAL (cumulative)** | 24.80 µL | | |

16. Once the RT program is finished, spin down the plate and add PCR master mix on ice/cold rack. Spin down the plate at 1000 g for 15 seconds. Take outside of the clean room workspace, place in a thermocycler and run the following program:

| Temperature | Time | Cycles |
|---|---|---|
| 98 C | 3 min | |
| 98 C | 00:15 | 22 cycles (single HSPCs) |
| 67 C | 00:20 | |
| 72 C | 6 min | |
| 72 C | 5 min | |
| 4 C | HOLD | |

**Bead clean-up – Timing: 45 minutes**

17. Add 16 µL of beads (Ampure XP Beads, Beckman Coulter, Cat. No. 391-02-501) into a V-shaped 96 well plate (AXYGEN, P-96-450V-C, 500 uL 96 well "V" bottom clear; Ref. 391-02-501).

18. Aliquot 11 µL of clean water (PCR grade) into the same V-shaped 96 well plate.

19. Aliquot 14 µL of each cDNA library into each well of the same plate and pipette up and down to mix cDNA with beads (0.6:1 beads to cDNA ratio). Incubate for 5 minutes at RT.

20. Incubate mixture on a 96-well magnetic stand for 2 minutes. Once the liquid is clear of beads, remove the liquid.

21. Wash the beads twice with 80 % EtOH (freshly prepared, dilute EtOH in PCR grade water). Add 100 µL of ethanol to each well, incubate for 30 seconds and remove. Repeat once more (2 times in total) and use P10 tips to remove any remaining ethanol.

22. Leave the beads to air-dry for 3 minutes. Be careful not to overdry the beads at this point or it will be difficult to resuspend them.

23. Resuspend the beads into 8 µL of EB Buffer (Qiagen Cat No./ID: 19086) with the plate off the magnet. Incubate for 30 seconds and put the plate back onto the magnet. Incubate into the magnet until the liquid is clear, then transfer 7.5 µL of purified cDNA library to a new plate for -20 C storage or further processing.

24. Check cDNA traces quality and size distributions using Bioanalyzer (Agilent), Fragment Analyzer Automated CE System (Advanced Analytical) or similar capillary system. Representative good quality cDNA traces are shown below (Figure 1).

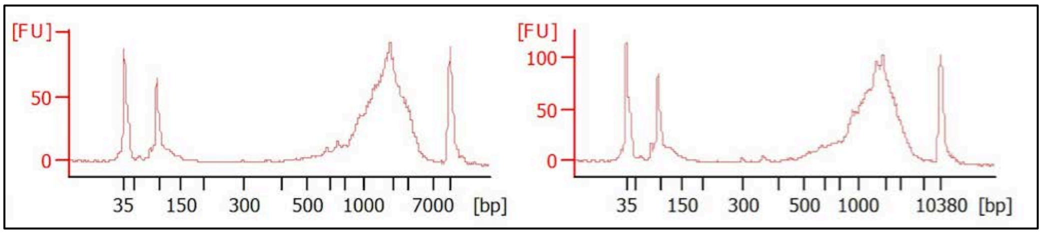


Figure 1. Representative bead-purified cDNA traces from single HSPCs synthesized using full-length TARGET-seq in 96-well plates.

**Whole transcriptome library preparation – Timing: 2 hours – 1.5 hours hands on time**

25. Library preparation is performed using a commercially available Nextera XT Kit (FC-131-1096, Illumina) and commercially available i5 and i7 indeces (Nextera XT Index

Kit, FC-131-1001, Illumina) using one fourth of the recommended volume. First, add 2.5 µL of Tagmentation Buffer into the required number of wells in a 96-well or 384-well plate (one well will be used for each cell).

26. Add 700 pg of purified cDNA from each pool in a total volume of 1.25 µL and 1.25 µL of Amplicon Tagmentation Mix (ATM). Incubate 6 minutes at 55 C (total volume 5 µL).

| **Reagents** | **1 reaction (µL)** |
|---|---|
| Tagmentation Buffer | 2.5 µL |
| Bead purified cDNA (560 pg/µL) | 1.25 µL |
| ATM (Amplicon Tagment Mix) | 1.25 µL |
| **TOTAL** | 5 µL |

27. Once the incubation is finished, add 1.25 µL of NT (Neutralization) buffer to neutralize the tagmentation reaction.

28. Prepare PCR master mix as outlined below.

| **Reagents** | **1 reaction (µL)** |
|---|---|
| i7 index (2 µM) | 1.25 µL |
| i5 index (2 µM) | 1.25 µL |
| NPM (PCR master mix) | 3.75 µL |
| **TOTAL** | 6.25 µL |
| **TOTAL (cumulative)** | 12.5 µL |

29. Incubate in a thermocycler and run the following PCR program:

| **Temperature** | **Time** | **Cycles** |
|---|---|---|
| 95 C | 30 seconds | 1 |
| 95 C | 10 seconds | 14 cycles |
| 55 C | 30 seconds | |
| 72 C | 30 seconds | |
| 72 C | 5 minutes | 1 |
| 4 C | HOLD | 1 |

30. Bead-purify libraries using Ampure XP beads. First, dilute the product 1:1 with 12.5 µL of PCR-grade water. Aliquot each barcoded and tagmented library into a V-shaped 96 well plate (Cat. No. P-96-450V-C, Axygen) and aliquot 16 µL of pre-warmed Ampure XP beads (Beckman Coulter; Cat. No. A63881) into each well (0.6:1 beads to cDNA ratio). Incubate for 5 minutes at RT.

31. Incubate mixture into a 96-well magnetic stand for 2 minutes. Once the liquid is clear of beads, remove the liquid.

32. Wash the beads twice with 80 % EtOH (freshly prepared, dilute EtOH in PCR grade water). Add 100 µL of ethanol to each well, incubate for 30 seconds and remove.

Repeat once more (2 times in total) and use P10 tips to remove any remaining ethanol.

33. Leave the beads to air-dry for 3 minutes. Be careful not to overdry the beads at this point or it will be difficult to resuspend them.

34. Resuspend the beads into 21 µL of EB Buffer (Qiagen Cat No./ID: 19086) with the plate off the magnet. Incubate for 30 seconds and put the plate back onto the magnet.

35. Incubate on the magnet until the liquid is clear of beads, then transfer 20 µL of purified tagmented/barcoded library to a new plate for -20 °C storage or further processing.

36. Run libraries on D5000 TapeStation or similar capillary array. Library fragments should be from 300 bp to 800 bp on average (Figure 2):

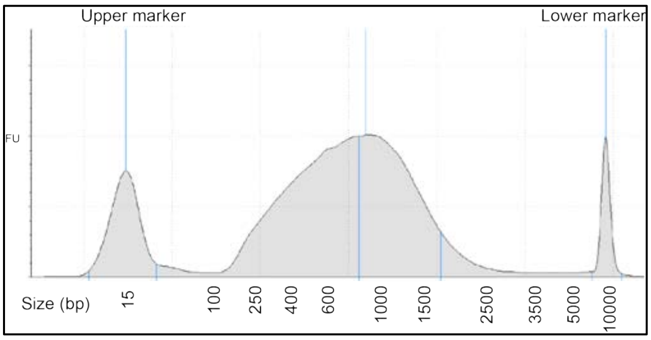

**Figure 2.** Representative traces of tagmented, amplified and bead-purified full-length Nextera XT libraries.

37. Quantify tagmented and barcoded libraries using Qubit (ThermoFisher, Cat. No. 32854) and pool equimolar concentrations of each library. Quantify the final pool and sequence on a NextSeq/HiSeq platform.

**Single cell genotyping library preparation for NGS – Timing: 5 hours – 2 hours hands on time**

38. Take one aliquot of the unpurified cDNA-amplicon mix, dilute 1:2 with PCR Grade water and use as an input for the first barcoding PCR (PCR1). Perform an individual PCR reaction for each sample in a 384 well-plate (FrameStar 384, Cat. No. 4ti-0384/C). During this PCR reaction, target-specific primers attached to universal barcodes (CS1/CS2 adaptors) will be added to each amplicon from each sample, in order to prepare a targeted sequencing library. Targets with similar amplification efficiencies might be amplified simultaneously in the same reaction for the same single cell. Note: gDNA and cDNA pre-amplified amplicons don't have a cell-specific barcode at this stage; therefore, amplicons corresponding to each cell should be kept in individual wells of the 384 well-plate, taking precautions to avoid cross-well contamination.

39. Prepare PCR Mix and aliquot in the 384 well-plate using a Biomek FxP Liquid Handler (Beckman Coulter) of similar liquid handling platform:

| PCR1 BARCODING with target-specific primers | 1 Reaction | Storage | Cat. No. |
|---|---|---|---|
| KAPA 2G Ready Mix | 3.125 µL | -20 °C | KAPA 2G Robust HS PCR Kit #KK5517 |
| Primer F1+R1 (20 µM) | 0.375 µL | -20 °C | Custom primers (Invitrogen) cartridge purification, resuspend in TE |
| Primer F2+R2 (20 µM) | 0.375 µL | -20 °C | |
| Primer F3+R3 (20 µM) | 0.375 µL | -20 °C | |
| Primer FX+RX… | … | … | |
| RT-PCR Grade Water | Variable | -20 °C | UltraPure DNAse/RNAse-Free Distilled Water, (Life Technologies, #10977035) |
| cDNA aliquot | 1.5 µL | -20 °C | |
| **TOTAL** | 6.25 µL | | |

40. Incubate in a thermocycler and run the following PCR program:

| PCR1 PROGRAM | | |
|---|---|---|
| Temperature | Time (min:sec) | Cycles |
| 95 C | 03:00 | 1 |
| 95 C | 00:20 | 20 |
| 60 C | 00:15 | |
| 72 C | 01:00 | |
| 72 C | 05:00 | 1 |
| 4 C | HOLD | |

41. Use 2.5 µL of PCR1 product as an input for the next reaction (PCR2). During this step, sample-specific barcodes are attached to previously barcoded amplicons using the Access Array™ Barcode Library for Illumina® Sequencers (384, Single Direction, Fluidigm). Barcode each sample in individual reactions.

42. Aliquot the barcodes (Access Array™ Barcode Library for Illumina® Sequencers) into a 384 well plate, and aliquot the PCR1 product into the same plate using a Biomek FxP Liquid Handler (Beckman Coulter) of similar liquid handling platform.

43. Prepare the PCR2 master mix and aliquot:

| PCR2 BARCODING with Illumina compatible primers | 1 Reaction | Storage | Cat. No. |
|---|---|---|---|
| FastStart High Fidelity 10X Reaction Buffer | 1 µL | -20 °C | FastStart High Fidelity PCR System REF:04738292001 |
| MgCl2 (25 mM) | 1.8 µL | -20 °C | |
| DMSO | 0.5 µL | -20 °C | |
| dNTP Mix (10 mM) | 0.2 µL | -20 °C | |
| FastStart High Fidelity Enzyme (5U/µL) | 0.1 µL | -20 °C | |

| | | | |
|---|---|---|---|
| RT-PCR Grade Water | 1.90 µL | -20 °C | UltraPure DNAse/RNAse-Free Distilled Water, (Life Technologies, #10977035) |
| Single-direction barcodes (2 µM, Fluidigm) | 2.0 µL | -20 °C | Access Array™ Barcode Library for Illumina® Sequencers-384, Single Direction, Fluidigm (Cat. No. 100-4876) |
| PCR1 barcoding aliquot | 2.5 µL | -20 °C | |
| **TOTAL** | 10 µL | | |

44. Incubate in a thermocycler and run the following PCR program:

| **PCR2 PROGRAM** | | |
|---|---|---|
| Temperature | Time (min:sec) | Cycles |
| 98 C | 10:00 | 1 |
| 98 C | 00:20 | 10 |
| 60 C | 00:15 | |
| 72 C | 01:00 | |
| 72 C | 03:00 | 1 |
| 4 C | HOLD | |

45. Pool amplicons from each barcoded library using a liquid handling platform and use Ampure XP beads to clean-up pooled libraries (0.8:1 beads to cDNA ratio). Quantify libraries using Qubit (ThermoFisher; Cat No. 32854) and check library size distribution and specific amplification of targeted amplicons on D1000 TapeStation or similar capillary array (Figure 3). Note: barcodes and adaptors add 103 bp extra to the original PCR product.

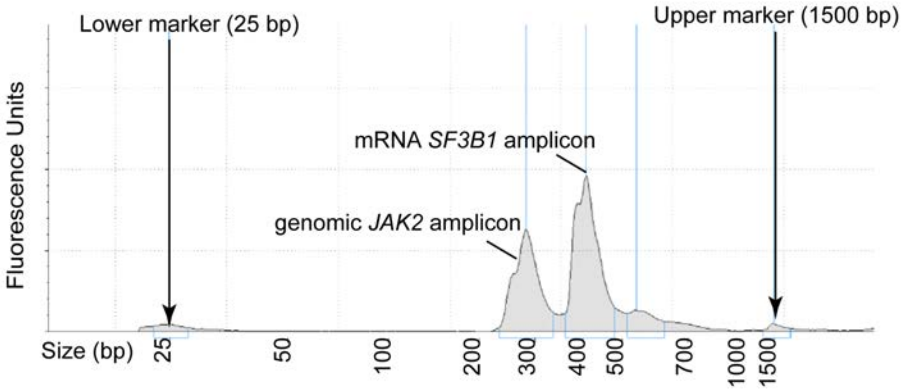


**Figure 3.** Representative distributions of targeted amplicon libraries from genomic *JAK2* and mRNA *SF3B1* amplicons in a multiplexed reaction.

46. Libraries are ready for sequencing using custom sequencing primers targeted to CS1/CS2 tags (500 nM of CS1 and CS2 primers in a total volume of 700 µL for R1 and R2; 500 nM of CS1rc and CS2rc primers in a total volume of 700 µL for Index Read when using the MiSeq platform, Illumina). Note: CS1/CS2 and CS1rc/CS2rc sequencing primers contain LNA modifications (see Key Resources), as compared to CS1/CS2 tags used for PCR1 target-specific primers.
